# *Listeria* phages induce Cas9 degradation to protect lysogenic genomes

**DOI:** 10.1101/787200

**Authors:** Beatriz A. Osuna, Shweta Karambelkar, Caroline Mahendra, Kathleen A. Christie, Bianca Garcia, Alan R. Davidson, Benjamin P. Kleinstiver, Samuel Kilcher, Joseph Bondy-Denomy

## Abstract

Bacterial CRISPR-Cas systems employ RNA-guided nucleases to destroy foreign DNA. Bacteriophages, in turn, have evolved diverse “anti-CRISPR” proteins (Acrs) to counteract acquired immunity. In *Listeria monocytogenes*, prophages encode 2-3 distinct anti-Cas9 proteins, with *acrIIA1* always present; however, its mechanism is unknown. Here, we report that AcrIIA1 binds with high affinity to Cas9 via the catalytic HNH domain and, in *Listeria*, triggers Cas9 degradation. AcrIIA1 displays broad-spectrum inhibition of Type II-A and II-C Cas9s, including an additional highly-diverged *Listeria* Cas9. During lytic infection, AcrIIA1 is insufficient for rapid Cas9 inactivation, thus phages require an additional “partner” Acr that rapidly blocks Cas9-DNA-binding. The AcrIIA1 N-terminal domain (AcrIIA1^NTD^) is dispensable for anti-CRISPR activity; instead it is required for optimal phage replication through direct transcriptional repression of the anti-CRISPR locus. AcrIIA1^NTD^ is widespread amongst *Firmicutes*, can repress anti-CRISPR deployment by other phages, and has been co-opted by hosts potentially as an “anti-anti-CRISPR.” In summary, *Listeria* phages utilize narrow-spectrum inhibitors of DNA binding to rapidly inactivate Cas9 in lytic growth and the broad-spectrum AcrIIA1 to stimulate Cas9 degradation for protection of the *Listeria* genome in lysogeny.

## INTRODUCTION

All cells must combat viral infections to survive. Bacteria have evolved innate and adaptive defense mechanisms against bacterial viruses (phages), which constantly pose a risk of infection. One such defense mechanism is CRISPR-Cas, a common and diverse adaptive immune system in prokaryotes that encompasses two distinct classes and six types (I-VI) (Koonin et al., 2017; Makarova et al., 2015). The CRISPR array maintains a genetic record of past viral infections with phage DNA fragments (spacers) retained between clustered regularly interspaced short palindromic repeats (CRISPR) (Mojica et al., 2005). These phage-derived spacers are transcribed into CRISPR RNAs (crRNAs) that complex with Cas nucleases to guide the sequence-specific destruction of invading nucleic acids (Brouns et al., 2008; Garneau et al., 2010). The CRISPR-associated (cas) genes typically neighbor the CRISPR array and encode proteins that facilitate spacer acquisition into the CRISPR array (Nuñez et al., 2014; Yosef et al., 2012), generate mature crRNAs (Deltcheva et al., 2011; Haurwitz et al., 2010), and cleave invading genomes (Garneau et al., 2010).

To counteract bacterial immunity, phages have evolved multiple mechanisms of CRISPR-Cas evasion (Borges et al., 2017). Phage-encoded anti-CRISPR proteins that directly inhibit type I-C, I-D, I-E, I-F, II-A, II-C, and V-A CRISPR-Cas systems have been identified (Hwang and Maxwell, 2019; Trasanidou et al., 2019). These anti-CRISPRs have distinct protein sequences, structures, and mechanisms of inactivation. Some anti-CRISPRs block CRISPR-Cas target DNA binding by steric occlusion and DNA mimicry (Bondy-Denomy et al., 2015; Dong et al., 2017; Jiang et al., 2019; Liu et al., 2019; Shin et al., 2017; Yang and Patel, 2017), guide-RNA loading interference (Thavalingam et al., 2019; Zhu et al., 2019), and effector dimerization (Fuchsbauer et al., 2019; Harrington et al., 2017; Zhu et al., 2019). Other anti-CRISPRs prevent DNA cleavage by interacting with the catalytic domain of Cas nucleases (Bondy-Denomy et al., 2015; Harrington et al., 2017). Anti-CRISPRs that inactivate Type II CRISPR-Cas systems, which are widely utilized for genome editing applications, have been extensively characterized in biochemical and heterologous cell-based systems (Bondy-Denomy, 2018; Yao et al., 2018). However, few studies have examined anti-CRISPR functions in the natural context of phage-bacteria warfare (Hynes et al., 2017, 2018).

In the lytic cycle, phage replication causes host cell lysis, whereas in lysogeny, temperate phages integrate into the bacterial chromosome and become prophages. The bacterial host and prophage replicate together during lysogeny and prophages can contribute novel genes that provide fitness benefits or even serve as regulatory switches (Argov et al., 2017; Bondy-Denomy et al., 2016; Borges et al., 2017; Chen et al., 2005; Feiner et al., 2015). In *Listeria monocytogenes*, some prophages employ “active lysogeny” during mammalian cell infection, wherein temporary prophage excision from the bacterial chromosome allows expression of the *comK* gene required for *Listeria* replication in macrophages (Rabinovich et al., 2012). Prophages also inactivate CRISPR-Cas in *L. monocytogenes* through the expression of anti-CRISPR proteins (Rauch et al., 2017). In lysogens with CRISPR arrays encoding spacers that target the prophage (i.e. self-targeting), anti-CRISPRs are essential for host and prophage survival. Whether anti-CRISPRs play distinct roles during lysogeny or lytic growth when expressed by temperate phages is unknown.

Here, we show that the *Listeria* phage protein AcrIIA1 selectively triggers degradation of catalytically active Cas9, through a direct interaction between the AcrIIA1^CTD^ (C-terminal domain) unstructured loop and Cas9 HNH domain. AcrIIA1 is sufficient to stabilize CRISPR-targeted prophages, but is ineffective during lytic replication. This inactivity necessitates the co-existence of AcrIIA1 with an anti-CRISPR (e.g. AcrIIA2, AcrIIA4, or AcrIIA12, identified here) that rapidly blocks Cas9 during lytic infection. While highly conserved across AcrIIA1 homologs, the AcrIIA1^NTD^ (N-terminal domain) is completely dispensable for anti-CRISPR activity and is instead a crucial repressor of *acr* locus transcription, a requirement for optimal phage fitness.

## RESULTS

### AcrIIA1 interacts with Cas9 and triggers its degradation

To determine the AcrIIA1 mechanism of action, we first attempted to immunoprecipitate Cas9 from *L. monocytogenes* (*Lmo*10403s) strains, where AcrIIA1 was expressed from one of three prophages (ΦA006, ΦA118, and ΦJ0161a). Surprisingly, upon immunoblotting for Cas9 protein, we observed highly reduced Cas9 levels in these lysogens (Figure 1A). Transcriptional and translational reporters revealed that transcript levels were unaffected, while the protein reporter levels decreased by ∼70% (Figure 1A). RT-qPCR experiments confirmed Cas9 mRNA levels were unaffected in each lysogen (Figure S1A). AcrIIA1 alone, but not AcrIIA4, was sufficient to mediate decreased Cas9 levels in both reporter and western blot assays (Figure 1B). The well-studied orthologue, SpyCas9 (53% amino acid identity to LmoCas9), displayed the same post-transcriptional AcrIIA1-dependent loss of Cas9 when introduced into *L. monocytogenes* (Figure 1B).

**Figure 1.**
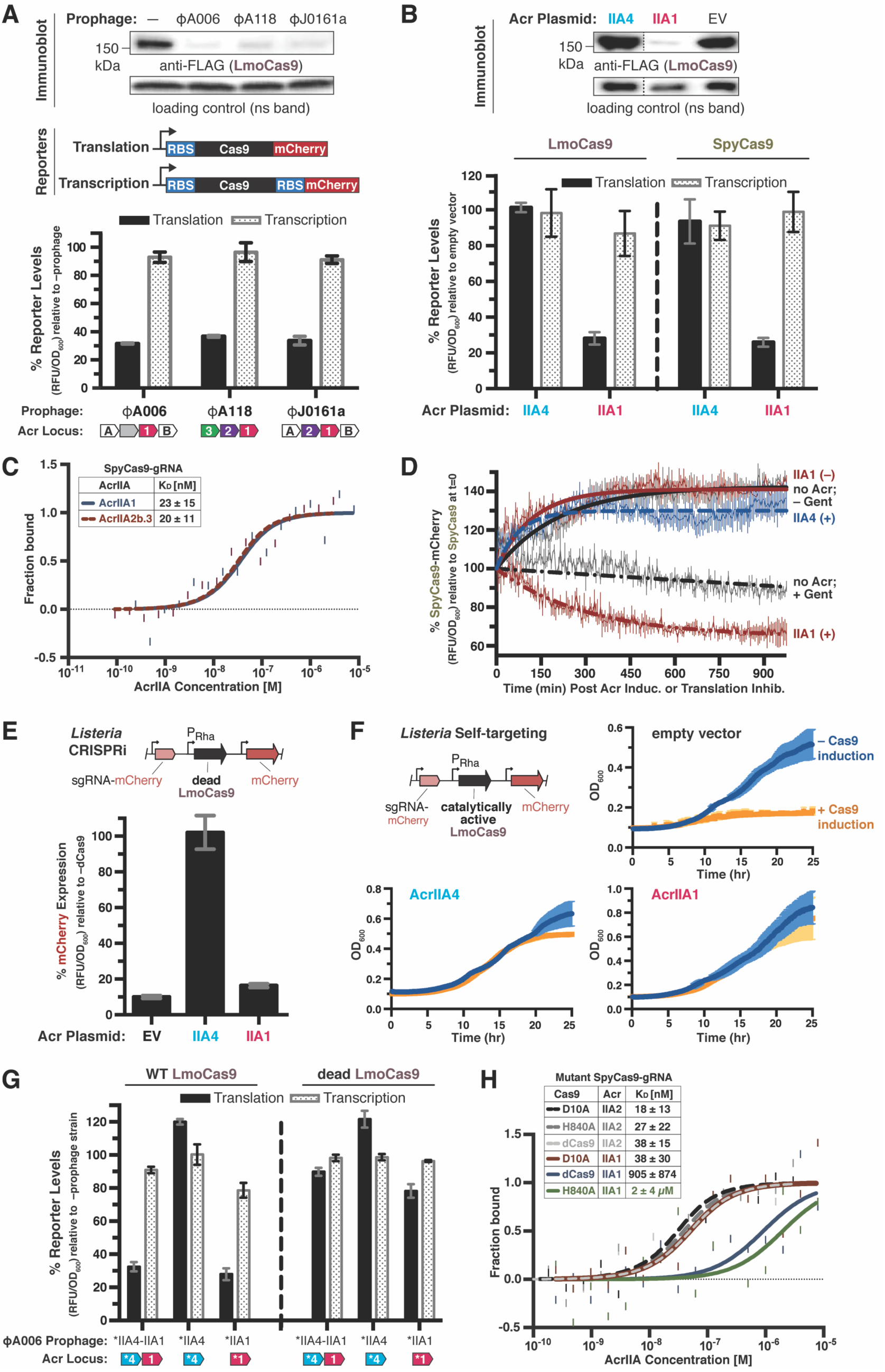
AcrIIA1 Binds Catalytically Active Cas9 and Triggers its Degradation in *Listeria*. (A and B) Immunoblots detecting FLAG-tagged LmoCas9 protein and a non-specific (ns) protein loading control in *Listeria monocytogenes* strain 10403s (*Lmo*10403s) lysogenized with the indicated wild-type prophages (A, top) or *Lmo*10403s containing Acr-expressing plasmids (B, top). Dashed lines indicate where intervening lanes were removed for clarity (B, top). Representative blots of at least three biological replicates are shown (A and B). Schematics of translational and transcriptional reporters used to measure Lmo or Spy Cas9 protein and mRNA levels in *Lmo*10403s (A, middle). Cas9 translational (black bars) and transcriptional (gray shaded bars) reporter measurements reflect the mean percentage mCherry relative fluorescence units (RFU normalized to OD_600_) in the indicated lysogens (A, bottom) or strains with Acr-expressing plasmids (B, bottom) relative to the control strain lacking a prophage (– prophage) (A, bottom) or containing an empty vector (B, bottom). Error bars represent the mean ± SD of at least three biological replicates. (C) Quantification of the binding affinities (K_D_; boxed inset) of AcrIIA1 and AcrIIA2b.3 for WT SpyCas9-gRNA using microscale thermophoresis. Data shown are representative of three independent experiments. (D) SpyCas9-mCherry protein levels post anti-CRISPR induction or translation inhibition. *Lmo*10403s strains expressing SpyCas9-mCherry from the constitutively active pHyper promoter and AcrIIA1 or AcrIIA4 from an inducible promoter were grown to mid-logarithmic phase and treated with 100 mM rhamnose to induce Acr expression (+, thick dashed lines) or 100 mM glycerol as a neutral carbon source control (–, thick solid lines) and 5 µg/mL gentamicin (Gent) to inhibit translation (+) or water (–) as a control. SpyCas9-mCherry protein measurements reflect the mean percentage fluorescence (RFUs normalized to OD_600_) relative to the SpyCas9-mCherry levels at the time (0 min) translation inhibition was initiated (thin solid lines). Error bars (vertical lines) represent the mean ± SD of at least three biological replicates. Data were fitted by nonlinear regression to generate best-fit decay curves (thick lines). See Figure S1D for additional controls and S1E for data showing tight repression of the pRhamnose promoter under non-inducing conditions. (E and F) Anti-CRISPR inhibition of CRISPRi (E) or self-targeting (F) in *Listeria*. *Lmo*10403s strains contain chromosomally-integrated constructs expressing dead (E) or catalytically active (F) LmoCas9 from the inducible pRha-promoter and a sgRNA that targets the pHelp-promoter driving mCherry expression. For CRISPRi, mCherry expression measurements reflect the mean percentage fluorescence (RFU normalized to OD_600_) in deadCas9-induced cells relative to uninduced (–dCas9) controls of three biological replicates ± SD (error bars) (E). For self-targeting, bacterial growth was monitored after LmoCas9 induction (orange lines) or no induction (blue lines) and data are displayed as the mean OD_600_ of three biological replicates ± SD (error bars) (F). See Figure S1F for CRISPRi data with *Lmo*10403s expressing dead SpyCas9. (G) Translational (black bars) and transcriptional (gray shaded bars) reporter levels of catalytically active (left) and dead LmoCas9 (right) in *Lmo*10403s lysogenized with engineered isogenic ΦA006 prophages encoding the indicated anti-CRISPRs. Measurements were normalized and graphed as in (A, bottom) with error bars representing the mean ± SD of at least three biological replicates. (*) Indicates the native orfA RBS (strong) in ΦA006 was used for Acr expression. See Figure S1H for equivalent data with *Lmo*10403s expressing SpyCas9. (H) Quantification of the binding affinities (K_D_; boxed inset) of AcrIIA1 (IIA1, solid lines) and AcrIIA2b.3 (IIA2, dashed lines) for catalytically dead (dCas9) and nickase (D10A or H840A) SpyCas9-gRNA complexes using microscale thermophoresis. Data shown are representative of three independent experiments.

To assay for a direct interaction *in vitro*, AcrIIA1 and SpyCas9 were purified (LmoCas9 was insoluble). AcrIIA1 and the SpyCas9-gRNA complex interacted with high affinity (K_D_ = 23 ± 15 nM) by microscale thermophoresis (MST), comparable to AcrIIA2b.3 (K_D_ = 20 ± 11 nM), a well-characterized Cas9-interactor (Jiang et al., 2019; Liu et al., 2019) (Figure 1C). Additionally, AcrIIA1 interacted with ApoCas9, unlike AcrIIA2b.3, suggesting a unique binding mechanism (Figure S1B). Neither binding event was sufficient to degrade Cas9 *in vitro*, nor was the protein destabilized when subjected to limited proteolysis (Figure S1C). We therefore considered whether the cellular environment of *L. monocytogenes* stimulates Cas9 degradation when bound by AcrIIA1. Indeed, we observed an accelerated decay of SpyCas9 protein upon induction of AcrIIA1 compared to treatment with a translation inhibitor, gentamicin (Figures 1D and S1D). In contrast, SpyCas9 protein increased over time when AcrIIA1 was not induced; similar to strains expressing AcrIIA4 or lacking an anti-CRISPR (Figures 1D and S1D). However, we paradoxically observed that AcrIIA1 did not inhibit catalytically-dead Cas9 (dCas9) in a CRISPRi assay using Lmo- or Spy-dCas9 engineered to repress RFP expression (Figures 1E and S1F), but did inhibit active Cas9 in an isogenic self-targeting strain (Figure 1F). Consistent with these findings, lysogens expressing AcrIIA1 or AcrIIA4 alone or together also revealed no significant decrease in dCas9 levels (Figures 1G and S1H), whereas active Cas9 protein diminished by ∼70% in all AcrIIA1-expressing lysogens (Figures 1G, S1G and S1H). Therefore, AcrIIA1 has a mechanism to detect catalytically active Cas9 protein and trigger its degradation.

Given the discrepant outcomes between Cas9 and dCas9, the ability of AcrIIA1 to bind these proteins *in vitro* was assessed. AcrIIA1 interacted with dCas9-gRNA ∼40-fold weaker (K_D_ = 905 ± 874 nM) than with Cas9-gRNA (Figures 1C, 1H and S1I). Only two residues differ between catalytically active Cas9 and dCas9 (D10A and H840A). AcrIIA1 binding to Cas9(D10A) (K_D_ = ∼38 nM) was similar to wild-type Cas9 (K_D_ = ∼23, Figure 1), whereas binding to Cas9(H840A) was ∼80-fold weaker (K_D_ = 2 ± 4 µM) (Figures 1H and S1I). AcrIIA2b.3, which binds the PAM-interacting domain, displayed no difference in binding affinity to the four Cas9 variants (K_D_ = 18 – 38 nM) (Figures 1H and S1I). Therefore, we conclude that AcrIIA1 triggers the degradation of catalytically active Cas9 in *L. monocytogenes* through a direct interaction with the Cas9 HNH domain (where H840 resides).

### AcrIIA1 protects CRISPR-targeted prophages but fails during lytic replication

Given that AcrIIA1 triggers Cas9 degradation, a mechanism not previously observed for any anti-CRISPR, we sought to determine when this activity manifests in the phage life cycle. Isogenic ΦA006 phages were engineered to encode no anti-CRISPR, *acrIIA1*, *acrIIA4*, or *acrIIA1* and *acrIIA4* together, and assessed along with wild-type (WT) phages, during lytic and lysogenic infection. When infecting *Lmo*10403s expressing Cas9 and a native ΦA006-targeting spacer sequence, phages encoding only *acrIIA1* surprisingly failed to replicate, similar to a Δ*acr* phage (efficiency of plaquing, EOP ≤ 3×10^−5^, Figures 2A and S2A). Phages encoding *acrIIA4* replicated well (EOP = 0.1 – 0.7, depending *on acrIIA4* expression strength), similar to WT ΦA006 (EOP ≥ 0.7), with no added benefit derived from *acrIIA1* (Figures 2A and S2A). In contrast, during lysogeny, ΦA006 prophages encoding *acrIIA1* completely prevented self-targeting upon Cas9 induction, whereas lysogens lacking an anti-CRISPR (Δ*acr*) died (Figure 2B). The remarkable difference in AcrIIA1 efficacy during lytic and lysogenic growth bolsters a conclusion that the Cas9 degradative mechanism is optimal for the lysogenic lifestyle, but not fast enough for inactivation during lytic replication.

**Figure 2.**
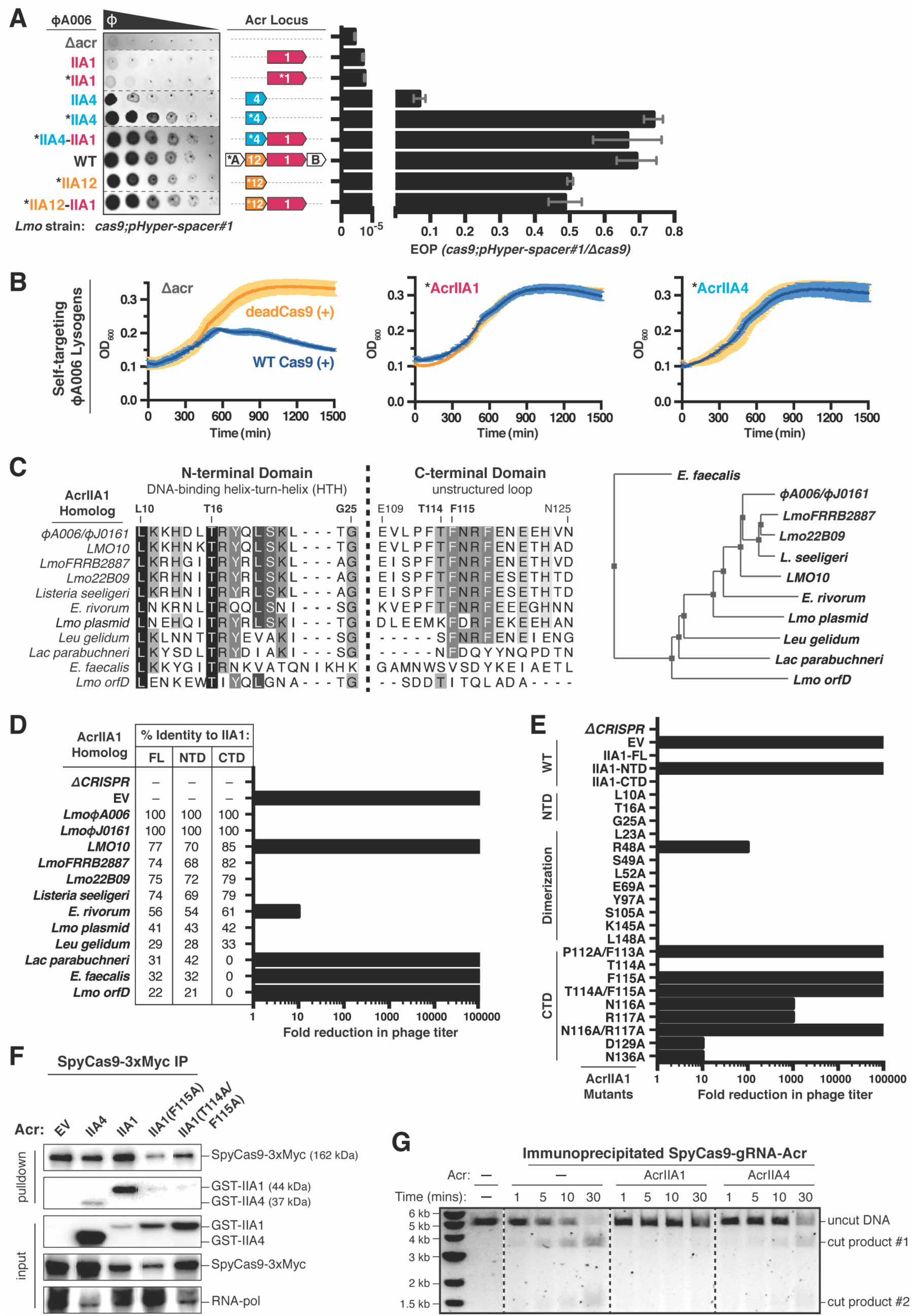
AcrIIA1 Inhibits Cas9 DNA Cleavage to Protect Prophages During Lysogeny. (A) Left: Representative image of plaquing assays where isogenic ΦA006 phages are titrated in ten-fold serial dilutions (black spots) on a lawn of *Lmo*10403s (gray background). Dashed lines indicate where intervening rows were removed for clarity. Right: Efficiency of plaquing (EOP) of isogenic ΦA006 phages (expressing the indicated anti-CRISPRs) on *Lmo*10403s. Plaque forming units (PFUs) were quantified on *Lmo*10403s overexpressing the first spacer in the native CRISPR array that targets ΦA006 (*cas9;pHyper-spacer#1*) and normalized to the number of PFUs measured on a non-targeting *Lmo*10403s-derived strain (Δ*cas9*). Data are displayed as the mean EOP of at least three biological replicates ± SD (error bars). See Figure S2A for EOP measurements of additional ΦA006 phages. (B) Bacterial growth curves of self-targeting *Lmo*10403s::ΦA006 isogenic lysogens expressing the indicated anti-CRISPRs and rhamnose-induced (+) WT or dead LmoCas9. WT LmoCas9 induction (blue lines), but not dead LmoCas9 (orange lines) is lethal in an Acr-deficient (Δacr) strain because the *Lmo*10403s CRISPR array contains a spacer targeting the ΦA006 prophage integrated in the bacterial genome. Data are displayed as the mean OD_600_ of at least three biological replicates ± SD (error bars) as a function of time (min). (*) Indicates the native orfA RBS (strong) in ΦA006 was used for Acr expression. (C) Left: Alignment of AcrIIA1 homolog protein sequences denoting key residues. Right: Phylogenetic tree of the protein sequences of AcrIIA1 homologs. See Figure S4D for a complete alignment of the AcrIIA1 homolog protein sequences. (D and E) Fold reduction in phage titer in response to SpyCas9 targeting of a *P. aeruginosa* DMS3m-like phage in the presence of AcrIIA1 homologs (D) or mutants (E). The percent protein sequence identities of each homolog to the full-length (FL) or domains (NTD or CTD) of AcrIIA1_ΦA006_ (IIA1) are listed in (D). The displayed fold reductions in phage titer were qualitatively determined by examining three biological replicates of each phage-plaquing experiment. See Figure S3A for representative pictures of the corresponding phage-plaquing experiments. (F) Immunoblots detecting GST-tagged anti-CRISPR proteins that co-immunoprecipitated with Myc-tagged SpyCas9 in a *P. aeruginosa* strain heterologously expressing Type II-A SpyCas9-gRNA and the indicated Acrs. For input samples, one-hundredth lysate volume was analyzed to verify tagged protein expression and RNA-polymerase was used as a loading control. Representative blots of at least three biological replicates are shown. See Figure S2F for the reciprocal GST-Acr pulldown. (G) Time courses of SpyCas9 DNA cleavage reactions in the presence of the indicated anti-CRISPRs conducted with SpyCas9-gRNA-Acr (or no Acr, –) complexes immunoprecipitated from *P. aeruginosa*. Dashed lines indicate where intervening lanes were removed for clarity. Data shown are representative of three independent experiments. See Figure S2H for reactions with AcrIIA1 mutants.

Given the inability of AcrIIA1 to inhibit Cas9 during lytic infection, phages may need additional Cas9 inhibitors. Indeed, in 119 *Listeria* prophage genomes analyzed, 77% encode *acrIIA1* with at least one additional *acrIIA* gene (i.e. *acrIIA2-A4*), 13% possess *acrIIA1* without a known *acrIIA* neighbor (including WT ΦA006), and 10% encode *orfD* (a distant *acrIIA1* orthologue), along with other uncharacterized ORFs (Rauch et al., 2017). The WT ΦA006 phage, which has *acrIIA1* and no other known *acr*, replicated far better (EOP ≥ 0.7) than a phage encoding *acrIIA1* alone, suggesting an additional Cas9 inhibitor in this phage (Figures 2A and S2A). Engineered phages encoding the gene adjacent to *acrIIA1* restored phage lytic replication (EOP ≥ 0.5) and revealed a new anti-CRISPR, AcrIIA12, which also inhibited Lmo (but not Spy) dCas9-based CRISPRi (Figures 2A, S2A and S2B). Notably, we observed the presence of *acrIIA12* in every *acr* locus previously reported to encode only *acrIIA1*, indicating that prophages do not encode *acrIIA1* alone. Therefore, *Listeria* prophages most commonly encode *acrIIA1*, which triggers Cas9 degradation to ensure stable lysogeny, in combination with a Cas9 interactor that blocks DNA binding (AcrIIA2, AcrIIA4, or AcrIIA12) for successful lytic replication.

### AcrIIA1 utilizes an unstructured C-terminal loop to inactivate Cas9

The AcrIIA1 crystal structure revealed a two-domain architecture, with a helix-turn-helix (HTH)-containing AcrIIA1^NTD^ similar to known transcriptional repressors and an extended AcrIIA1^CTD^ of unknown function (Ka et al., 2018). Surprisingly, the AcrIIA1^CTD^ was sufficient for anti-CRISPR function, protection from self-targeting, and triggering Cas9 protein degradation, while the AcrIIA1^NTD^ displayed no evidence of Cas9 regulation (Figures S2C and S2D). To identify AcrIIA1 residues required for anti-CRISPR function, we conducted multi-sequence alignments and used our previously developed heterologous *P. aeruginosa* anti-SpyCas9 screening platform (Jiang et al., 2019). AcrIIA1 homologs were identified in mobile genetic elements of *Listeria, Enterococcus, Lactobacillus*, and *Leuconostoc* species, ranging from 22% to 77% protein sequence identity (Figures 2C and S4D). Homology was driven by obvious sequence similarity in the NTD, with CTD conservation in only a subset of proteins. AcrIIA1 homologs with conserved CTDs displayed anti-SpyCas9 activity (except AcrIIA1_LMO10_), whereas the three proteins with highly diverged CTDs (including *orfD*) did not (Figures 2D and S3A). Alanine scanning mutagenesis of the conserved amino acids present in AcrIIA1 homologs identified a stretch of aromatic and charged residues in an unstructured region of the AcrIIA1^CTD^ (P112 to R117) that were required for complete anti-CRISPR activity (Figures 2E and S3A). Expression levels of each mutant protein were unperturbed relative to WT AcrIIA1 (Figure S3B). The F115A mutation completely abolished anti-CRISPR function (Figures 2E and S3A) and the interaction with Cas9 (Figures 2F and S2F-G). In *Listeria*, AcrIIA1(F115A) and AcrIIA1(T114A/F115A) mutants failed to protect cells from genomic self-targeting (Figure S2C) and these mutations either completely (T114A/F115A) or partially (F115A) restored Cas9 protein levels (Figure S2E).

When verifying expression of AcrIIA1 mutants, we observed that AcrIIA1-mediated inhibition does not trigger Cas9 degradation in *P. aeruginosa* (Figure S3B). Yet, similar to in *Listeria*, AcrIIA1 still displayed robust anti-CRISPR activity, inactivating Cas9 in phage-targeting and self-targeting experiments, while not interfering with CRISPRi (Figures S3A and S3C). Since AcrIIA1 can apparently inhibit Cas9 without causing degradation, we immunoprecipitated Cas9 bound to AcrIIA1 or the control AcrIIA4 from *P. aeruginosa* (Figures 2F and S2F) and assessed DNA cleavage activity *in vitro*. Cas9 was functional when immunoprecipitated alone but inhibited when co-purified with AcrIIA1 or AcrIIA4 (Figure 2G and S2H). The AcrIIA1 mutants (F115A and T114A/F115A) interacted with Cas9 very weakly (Figures 2F, S2F-G) and had little impact on DNA cleavage (Figure S2H). Interestingly, *in vitro* experiments with individually purified proteins revealed that AcrIIA1 is not sufficient to inhibit Cas9-mediated DNA cleavage (Figure S2I), despite its strong binding affinity, suggesting an additional cellular factor is required to inactivate Cas9. This putative multi-step process may explain why inhibition does not manifest immediately during lytic growth. Therefore, AcrIIA1 utilizes conserved residues in its CTD to interact with the Cas9 HNH domain, blocking DNA cleavage and triggering Cas9 protein degradation in *Listeria*. In a foreign host potentially lacking the Cas9-degrading pathway, DNA cleavage inhibition manifests.

### AcrIIA1 is a broad-spectrum Cas9 inhibitor

Given the ability of AcrIIA1 to inactivate Cas9 via recognition of a highly conserved catalytic residue, we assessed inhibition of diverged Cas9 orthologues. In *Escherichia coli* strains expressing Type II-A, II-B, and II-C Cas9 proteins (Figure 3A) targeting phage Mu, AcrIIA1 intermediately or completely inhibited four Type II-C (Boe, Hpa, Cje, and Geo) and two Type II-A (Sau and Spy) Cas9s (Figures 3B and S3D). In contrast, AcrIIA2 only weakly inhibited Hpa and SpyCas9, while AcrIIA4 only inactivated SpyCas9 (Figure 3B). Considering the biological driver of broad-spectrum Cas9 inhibition by AcrIIA1, a smaller Type II-A Cas9 (1,078 a.a.) was recently discovered in *L. ivanovii* (LivCas9) (Hupfeld et al., 2018) with similarities to other small Cas9 proteins (e.g. SauCas9) and Type II-C orthologues (Figure 3A). In *Listeria* strains expressing the small LivCas9 variant programmed to target phage ΦP35 or ΦA511, AcrIIA1 inhibited LivCas9 (Figures 3C and S3E). Thus, AcrIIA1 displays broad-spectrum activity against diverged Cas9 nucleases, whereas the well-characterized DNA binding inhibitors, AcrIIA2 and AcrIIA4, are much narrower in their inhibitory spectrum. This broad-spectrum inhibition also likely explains the utility of AcrIIA1 to phages infecting *Listeria*, where two distinct Cas9 orthologues are encountered.

**Figure 3.**
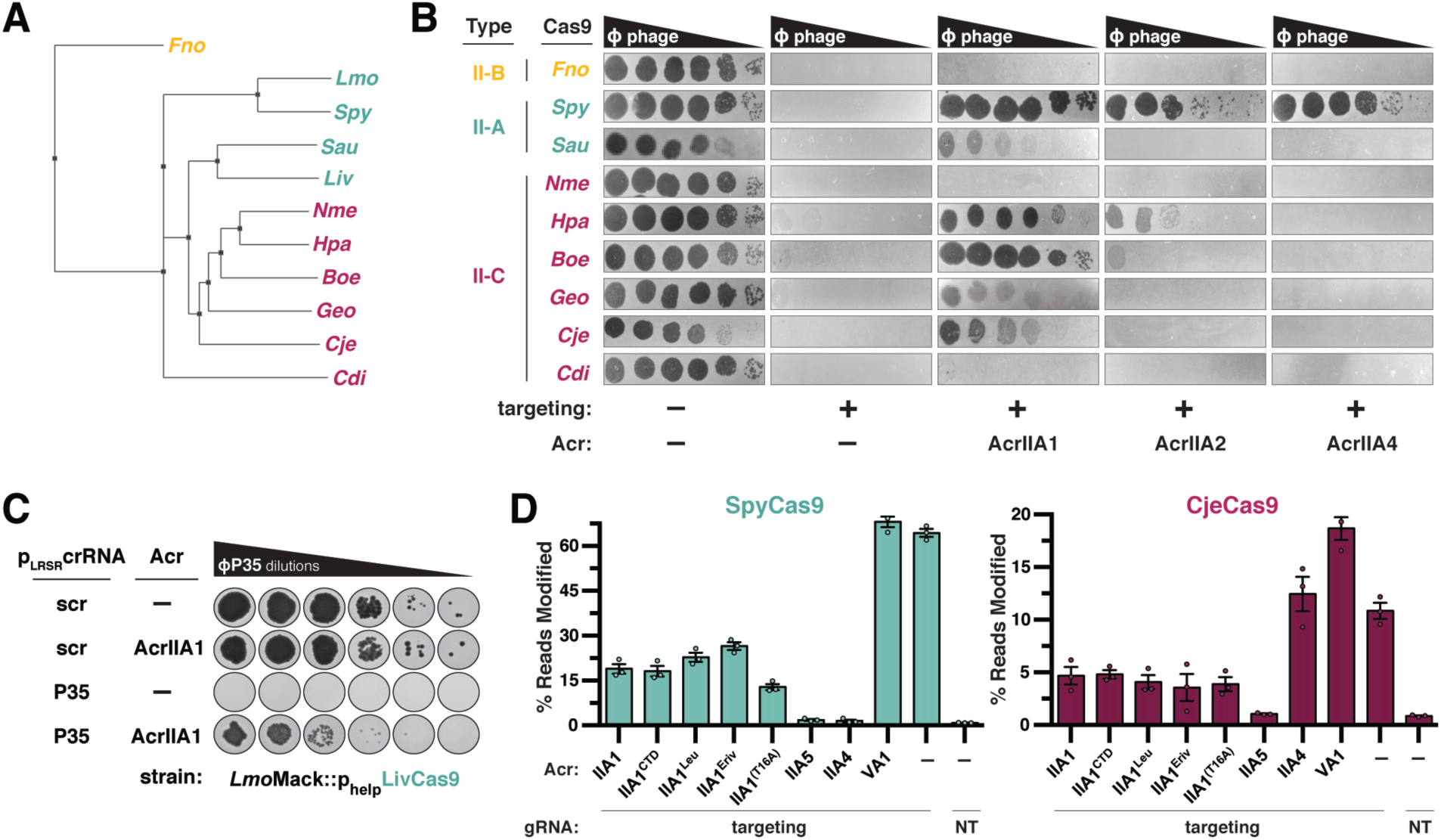
AcrIIA1 is a Broad-Spectrum Cas9 Inhibitor. (A) Phylogenetic tree of the protein sequences of Cas9 orthologues: Francisella novicida (Fno), Listeria monocytogenes (Lmo), Streptococcus pyogenes (Spy), Staphylococcus aureus (Sau), Listeria ivanovii (Liv), Neisseria meningitidis (Nme), Haemophilus parainfluenzae (Hpa), Brackiella oedipodis (Boe), Geobacillus stearothermophilus (Geo), Campylobacter jejuni (Cje), Corynebacterium diphtheriae (Cdi). (B) Plaquing assays where the *E. coli* phage Mu is titrated in ten-fold serial dilutions (black spots) on lawns of *E. coli* (gray background) expressing the indicated anti-CRISPR proteins and Type II-A, II-B and II-C Cas9-sgRNA combinations programmed to target phage DNA. Representative pictures of at least 3 biological replicates are shown. (C) Plaquing assays where the *Listeria* phage ΦP35 is titrated in ten-fold serial dilutions (black spots) on lawns of *L. monocytogenes* Mack (gray background) expressing episomal AcrIIA1 or no Acr (–), chromosomally-integrated LivCas9/tracrRNA, and episomal (pLRSR) crRNA that targets ΦP35 phage DNA or a non-targeting control (scr). (D) Gene editing activities of Cas9 orthologues in human cells in the presence of AcrIIA1 variants and orthologues. AcrIIA4 is a known inhibitor of SpyCas9, AcrIIA5 is a broad-spectrum inhibitor (Garcia et al., 2019, *in revision*), and AcrVA1 as a known non-inhibiting control for SpyCas9 orthologues. NT, no-sgRNA control condition. Error bars indicate SEM for three independent biological replicates. See Figure S3F for editing experiments with additional Cas9 orthologues.

The robust AcrIIA1 activity observed in various heterologous hosts led us to assess inhibition of Cas9 gene editing in human cells. We employed a deep sequencing-based approach to improve the dynamic range of edit detection, in comparison to our previous GFP-disruption assay (Rauch et al., 2017). HEK 293T cells were co-transfected with plasmids encoding *acrIIA1, cas9*, and sgRNAs targeting endogenous human sequences and editing efficacy was evaluated after 3 days. AcrIIA1 blocked the gene editing activity of SpyCas9 by 50-70% and of CjeCas9, SauCas9, St3Cas9, and NmeCas9 moderately, whereas AcrIIA4 only inhibited SpyCas9 (Figures 3D and S3E). Thus, AcrIIA1 inactivates diverse Cas9 orthologues in many heterologous systems, including bacteria (*L. monocytogenes, P. aeruginosa, E. coli*), yeast (Nakamura et al., 2019), and human cells, providing a genome editing modulator that specifically prevents Cas9 DNA cleavage. Future work is needed to enhance its efficiency, however.

### *acr* locus repression by AcrIIA1^NTD^ promotes general lytic growth and prophage induction

While interrogating the requirements for anti-CRISPR function, we observed that two engineered phages with deletions in their anti-CRISPR locus (ΦA006Δ*acr* and ΦJ0161aΔ*acrIIA1-2*) displayed a Cas9-independent lytic growth defect (Figure 4A). This defect was rescued by the provision of *acrIIA1^NTD^* in *trans* or by engineering an ΦA006 phage to express only the *acrIIA1^NTD^* (Figure 4A). Moreover, all engineered ΦA006 phages expressing an anti-CRISPR (e.g. *acrIIA1^CTD^*, *acrIIA4, acrIIA12*) without the *acrIIA1^NTD^* displayed a decrease in phage titer (PFU/mL) that was restored by *acrIIA1^NTD^ trans-* or *cis-*complementation (Figure 4A). The phage expressing only *acrIIA1^CTD^* (only observed fused to *acrIIA1^NTD^* in genomes) displayed the strongest lytic defect amongst the ΦA006 phages, while simply separating the two AcrIIA1 domains had no deleterious effect (Figure 4A). ΦJ0161aΔ*acrIIA1-2* had the most drastic lytic defect, failing to replicate unless complemented in *trans* with the *acrIIA1^NTD^* (Figure 4A). Moreover, the ΦJ0161aΔ*acrIIA1-2* prophage displayed a Cas9-independent prophage induction deficiency, yielding 25-fold less phage during mitomycin C induction, compared to the WT prophage or the *acrIIA1*-complemented mutant (Figure 4B). Attempts to efficiently induce ΦA006 prophages were unsuccessful, as previously observed (Loessner, 1991; Loessner et al., 1991). Therefore, aside from acting as an anti-CRISPR, AcrIIA1 plays an important Cas9-independent role in the phage life cycle, promoting optimal lytic replication and lysogenic induction.

**Figure 4.**
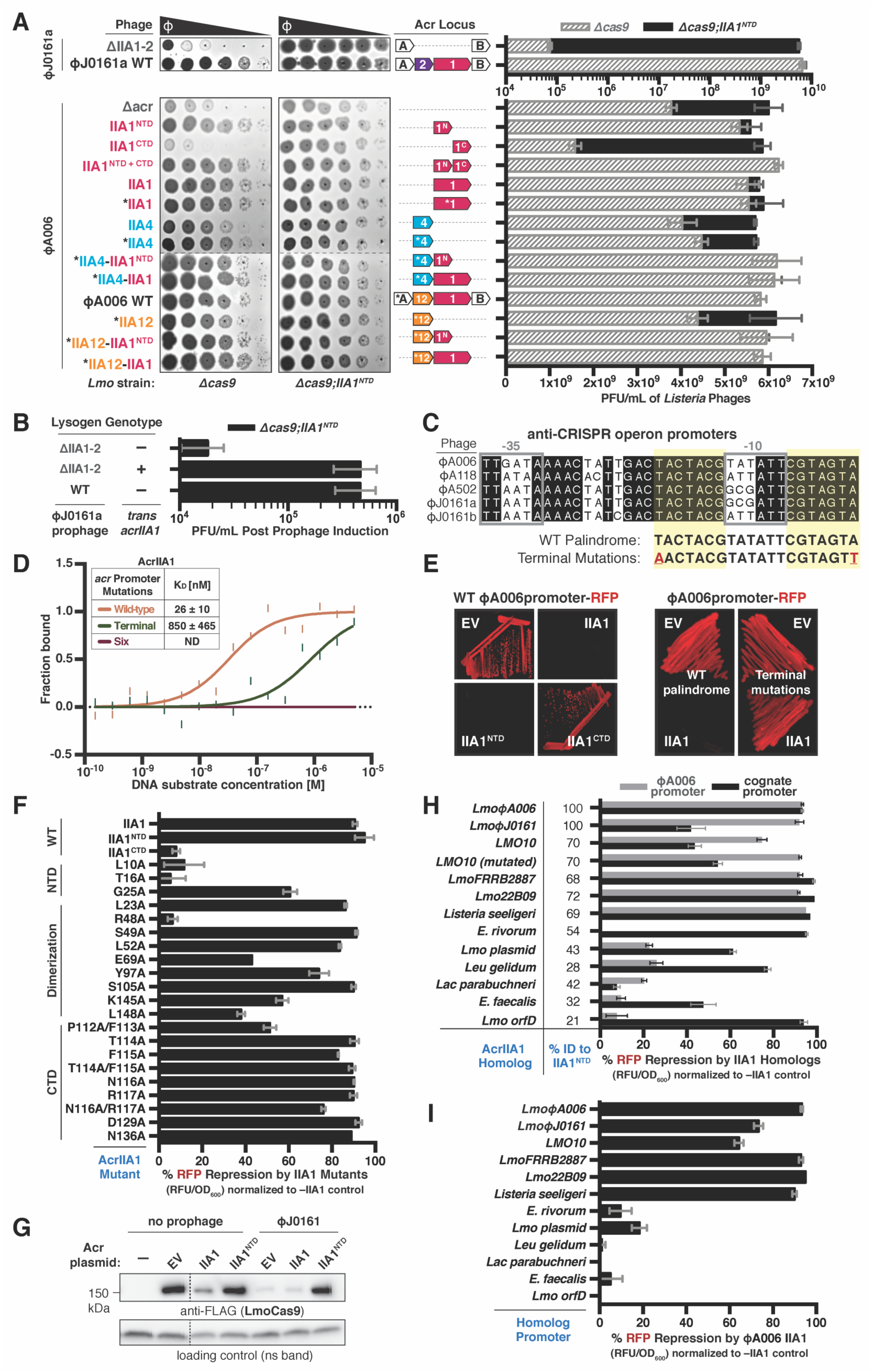
The AcrIIA1 N-terminal Domain Autorepresses the Anti-CRISPR Promoter. (A) Left: Representative images of plaquing assays where the indicated *Listeria* phages were titrated in ten-fold serial dilutions (black spots) on lawns of *Lmo*10403s (gray background) lacking Cas9 and encoding AcrIIA1^NTD^ (Δ*cas9;IIA1^NTD^*) or not (Δ*cas9*). Dashed lines indicate where intervening rows were removed for clarity. Right: Cas9-independent replication of isogenic ΦJ0161a or ΦA006 phages in *Listeria*. Plaque forming units (PFUs) were quantified on *Lmo*10403s lacking *cas9* (Δ*cas9*) and expressing AcrIIA1^NTD^ (black bars) or not (gray shaded bars). Data are displayed as the mean PFU/mL of at least three biological replicates ± SD (error bars). (B) Induction efficiency of ΦJ0161 prophages. Prophages were induced with mitomycin C from *Lmo*10403s::ΦJ0161 lysogens expressing *cis-acrIIA1* from the prophage Acr locus (WT) or not (ΔIIA1-2) and *trans-acrIIA1* from the bacterial host genome (+) or not (–). Plaque forming units (PFUs) were quantified on *Lmo*10403s lacking *cas9* and expressing AcrIIA1^NTD^ (Δ*cas9;IIA1^NTD^*). Data are displayed as the mean PFU/mL after prophage induction of four biological replicates ± SD (error bars). (C) Alignment of the phage anti-CRISPR promoter nucleotide sequences denoting the −35 and −10 elements (gray boxes) and conserved palindromic sequence (yellow highlight). Terminal palindrome mutations (red letters) were introduced for the binding assay in (D). See Figure S4A for a complete alignment of the promoters. (D) Quantification of the binding affinity (K_D_; boxed inset) of AcrIIA1 for the palindromic sequence within the *acr* promoter using microscale thermophoresis. ND indicates no binding detected. Data shown are representative of three independent experiments. (E) Expression of RFP transcriptional reporters containing the wild-type (left) or mutated (right) ΦA006-Acr-promoter in the presence of AcrIIA1 (IIA1) or each domain (IIA1^NTD^ or IIA1^CTD^). Representative images of three biological replicates are shown. (F and H-I) Repression of RFP transcriptional reporters containing the ΦA006-Acr-promoter (black bars in F; gray bars in H) or the cognate-AcrIIA1-homolog-promoters (black bars in H and I) by AcrIIA1_ΦA006_ (mutants in F; wild-type in I) or AcrIIA1 homolog (H) proteins. Data are shown as the mean percentage RFP repression in the presence of the indicated AcrIIA1 variants relative to controls lacking AcrIIA1 of at least three biological replicates ± SD (error bars). The percent protein sequence identities of each homolog to the ΦA006 AcrIIA1^NTD^ are listed in (H). (G) Immunoblots detecting FLAG-tagged LmoCas9 protein and a non-specific (ns) protein loading control in *Lmo*10403s::ΦJ0161a lysogens or non-lyosgenic strains containing plasmids expressing AcrIIA1 (IIA1) or AcrIIA1^NTD^ (IIA1^NTD^). Dashed lines indicate where intervening lanes were removed for clarity. Representative blots of at least three biological replicates are shown.

AcrIIA1^NTD^ contains an HTH motif with strong similarity to transcriptional repressors (Ka et al., 2018). Due to the Cas9-independent growth defects described above, we considered whether regulation of the anti-CRISPR locus is required. Alignments of the anti-CRISPR promoters of ΦA006, ΦJ0161, and ΦA118 revealed a highly conserved palindromic sequence (Figures 4C and S4A). An RFP transcriptional reporter assay showed that full-length AcrIIA1 and AcrIIA1^NTD^, but not AcrIIA1^CTD^, repress the ΦA006 anti-CRISPR promoter (Figure 4E, left). *In vitro* MST binding assays confirmed that AcrIIA1 (K_D_ = 26 ± 10 nM) or AcrIIA1^NTD^ (K_D_ = 28 ± 3 nM) bind the anti-CRISPR promoter with high affinity (Figures 4D and S4B). Moreover, mutagenesis of the palindromic sequence prevented AcrIIA1-mediated repression of the ΦA006 anti-CRISPR promoter (Figure 4E, right) and abolished promoter binding *in vitro* (Figure 4D). Alanine scanning mutagenesis of conserved residues predicted to be important for DNA binding and dimerization (Ka et al., 2018) identified AcrIIA1^NTD^ residues L10, T16, and R48 as critical for transcriptional repression, whereas AcrIIA1^CTD^ mutations had little effect (Figure 4F). Finally, we observed that Cas9 degradation induced by prophage-expressed AcrIIA1 in *L. monocytogenes* (Figure 1A) could be prevented by AcrIIA1^NTD^ overexpression, due to repression of the anti-CRISPR locus (Figure 4G). Thus, the AcrIIA1^NTD^-HTH domain represses anti-CRISPR transcription through a highly conserved operator, which is required for optimal phage fitness.

### Transcriptional autoregulation is a general feature of the AcrIIA1 superfamily

Recent studies have reported transcriptional autoregulation of anti-CRISPR loci by HTH-proteins in phages that infect Gram-negative *Proteobacteria*, as a mechanism to limit excessive transcription and downstream transcriptional conflict (Birkholz et al., 2019; Stanley et al., 2019). To determine whether anti-CRISPR locus regulation is similarly pervasive amongst mobile genetic elements in the Gram-positive *Firmicutes* phylum, we assessed AcrIIA1 homologs for transcriptional repression of their predicted cognate promoters and our model ΦA006 phage promoter. Homologs sharing amino acid sequence identity from 21% (i.e. OrfD) to 72% with AcrIIA1^NTD^ were selected from *Listeria, Enterococcus, Leuconostoc*, and *Lactobacillus* (Figure 4H and S4D). All AcrIIA1 homologs repressed transcription of their cognate promoters by 42-99%, except AcrIIA1 from *Lactobacillus parabuchneri*, where promoter expression was undetectable in a foreign host (Figures 4H and S4C). Strong repression of the model ΦA006 promoter was seen by *Listeria* orthologues possessing ≥68% protein sequence identity (Figure 4H). Likewise, AcrIIA1_ΦA006_ repressed the promoters of AcrIIA1 orthologues that repressed the ΦA006 promoter (Figure 4I). Interestingly, the AcrIIA1_LMO10_ homolog, which previously displayed no anti-CRISPR activity despite possessing 85% AcrIIA1^CTD^ sequence identity (Figures 2D and S3A), contains an AcrIIA1^NTD^ palindromic binding site overlapping its protein-coding sequence. AcrIIA1_LMO10_ anti-CRISPR function manifested when the AcrIIA1^NTD^ binding site was disrupted with silent mutations (Figure S3A). Altogether, these findings demonstrate that the anti-CRISPR promoter-AcrIIA1^NTD^ repressor relationship is highly conserved.

### Host-encoded AcrIIA1^NTD^ blocks phage anti-CRISPR deployment

Given that the AcrIIA1^NTD^ represses anti-CRISPR transcription, we wondered whether bacteria could co-opt this activity and manifest it in *trans*, inhibiting a phage from deploying its anti-CRISPR arsenal. We observed that ΦA006-derived phages encoding anti-CRISPRs were rendered vulnerable to Cas9 targeting when the host expressed anti-CRISPR-deficient AcrIIA1 A panel of distinct anti-CRISPR-encoding phages also became vulnerable to Cas9 targeting when AcrIIA1^NTD^ was expressed from a plasmid (Figure 5B) or from an integrated single-copy *acrIIA1^NTD^* driven by a prophage promoter (Figure S5A). Each of these phages possesses complete or partial spacer matches to the *Lmo*10403s CRISPR array. In contrast, replication of the non-targeted phage, ΦJ0161a, was unperturbed (Figure 5B). This demonstrates that host or mobile elements can use this repressor as an “anti-anti-CRISPR” to block anti-CRISPR synthesis, which may be particularly advantageous, if infecting phages encode other anti-CRISPR proteins (e.g. against the *Listeria* Type I-B CRISPR-Cas system).

**Figure 5.**
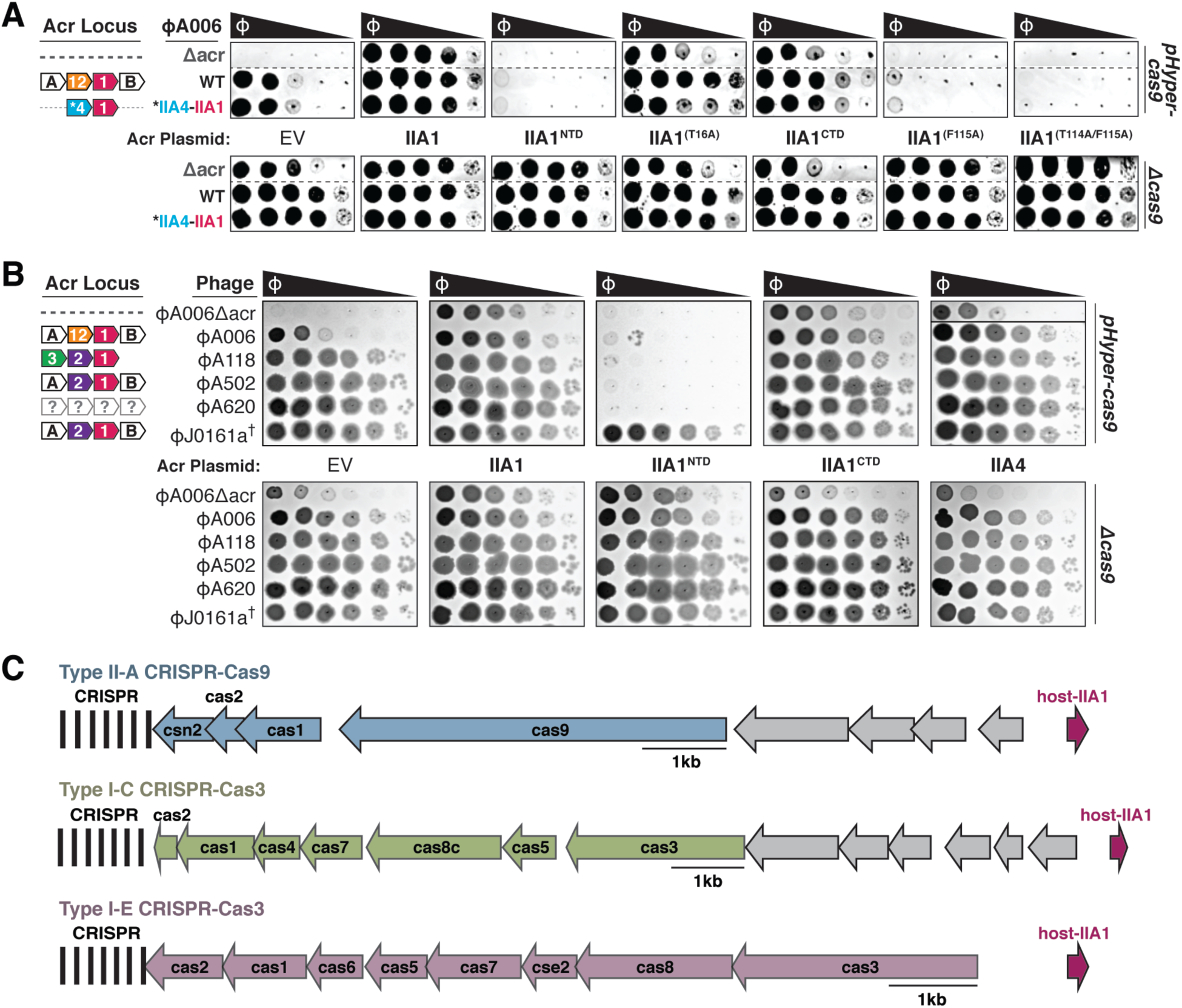
AcrIIA1^NTD^ Encoded from a Bacterial Host Displays “anti-anti-CRISPR” Activity. (A-B) Plaquing assays where engineered (A) or wild-type (B) *L. monocytogenes* phages are titrated in ten-fold dilutions (black spots) on lawns of *L. monocytogenes* (gray background) expressing anti-CRISPRs from plasmids, LmoCas9 from a strong promoter (*pHyper-cas9*) or lacking Cas9 (Δ*cas)*, and the natural CRISPR array containing spacers with complete or partial matches to the DNA of each phage. (†) Denotes the absence of a spacer targeting the ΦJ0161a phage. Representative pictures of at least 3 biological replicates are shown. Dashed lines indicate where intervening rows were removed for clarity (A). Solid lines indicate where separate images are shown. (C) Schematic of bacterial (host) AcrIIA1^NTD^ homologs encoded next to Type II-A, I-C, and I-E CRISPR-Cas loci in *Lactobacillus delbrueckii* strains.

The widespread prevalence of AcrIIA1 is driven by AcrIIA1^NTD^, with orthologues in many *Firmicutes* including *Enterococcus, Bacillus, Clostridium, and Streptococcus*. The AcrIIA1^NTD^ can be found either without a CTD or with a distinct CTD sequence. Diverged AcrIIA1^CTDs^ may represent novel anti-CRISPRs, inhibiting CRISPR-Cas systems in their respective hosts. In *Lactobacillus* sp., for example, there are full-length prophage proteins that lacked anti-SpyCas9 function and contain a novel AcrIIA1^CTD^ (Figures 2C, 2D and S3A). In other instances, core bacterial genomes encode AcrIIA1^NTD^ orthologues that are short ∼70-80 amino acid proteins possessing only the HTH domain. In particular, *Lactobacillus delbrueckii* strains contain an AcrIIA1^NTD^ homolog (35% identical, 62% similar to AcrIIA1_ΦA006_) with key residues conserved (e.g. L10 and T16). Although there are no known *Lactobacillus* phages that express anti-CRISPRs, this bacterial *acrIIA1^NTD^* gene may perform an “anti-anti-CRISPR” function. Remarkably, we observe that this AcrIIA1^NTD^ homolog is always a genomic neighbor of either the Type I-E, I-C, or II-A CRISPR-Cas systems in *L. delbrueckii* (Figure 5C). This association is supportive of a role that enables these CRISPR-Cas systems to function by repressing the deployment of phage inhibitors against each system. The functions of these diverse AcrIIA1 orthologues found in different bacteria, many of which act as transcriptional repressors (Figure 4H), remain to be elucidated.

## DISCUSSION

*Listeria* temperate phages commonly encode the multifunctional AcrIIA1 protein for protection against CRISPR-Cas and autorepression of anti-CRISPR transcription. The broad-spectrum AcrIIA1 is sufficient for Cas9 inactivation during lysogeny, but a nonfunctional anti-CRISPR during lytic growth, perhaps due to slow kinetics of Cas9 cleavage inhibition or degradation. Thus, AcrIIA1 always coexists with a distinct anti-Cas9 protein (e.g. AcrIIA2, AcrIIA4, AcrIIA12) that is much narrower in its inhibitory spectrum, but rapidly inactivates Cas9 during lytic replication. Therefore, *Listeria* temperate phages have evolved multiple anti-CRISPRs with distinct Cas9 binding sites and inactivation mechanisms because they synergistically grant unique advantages in each stage of the temperate phage life cycle (see model, Figure 6). While “partner” proteins AcrIIA4 and AcrIIA12 also protected CRISPR-targeted prophages, only AcrIIA1 triggered Cas9 degradation, presumably enhancing the likelihood of long-term stability in lysogeny. *Listeria* lysogens were devoid of Cas9 even when *acrIIA1* was co-encoded with other *acrs*, supporting that Cas9 degradation is the dominant inactivation mechanism in lysogeny. Given that Cas9 is required for selection of functional spacers by recognizing the correct PAM (Heler et al., 2015), eliminating this nuclease could also prevent acquisition of lethal self-targeting spacers.

**Figure 6.**
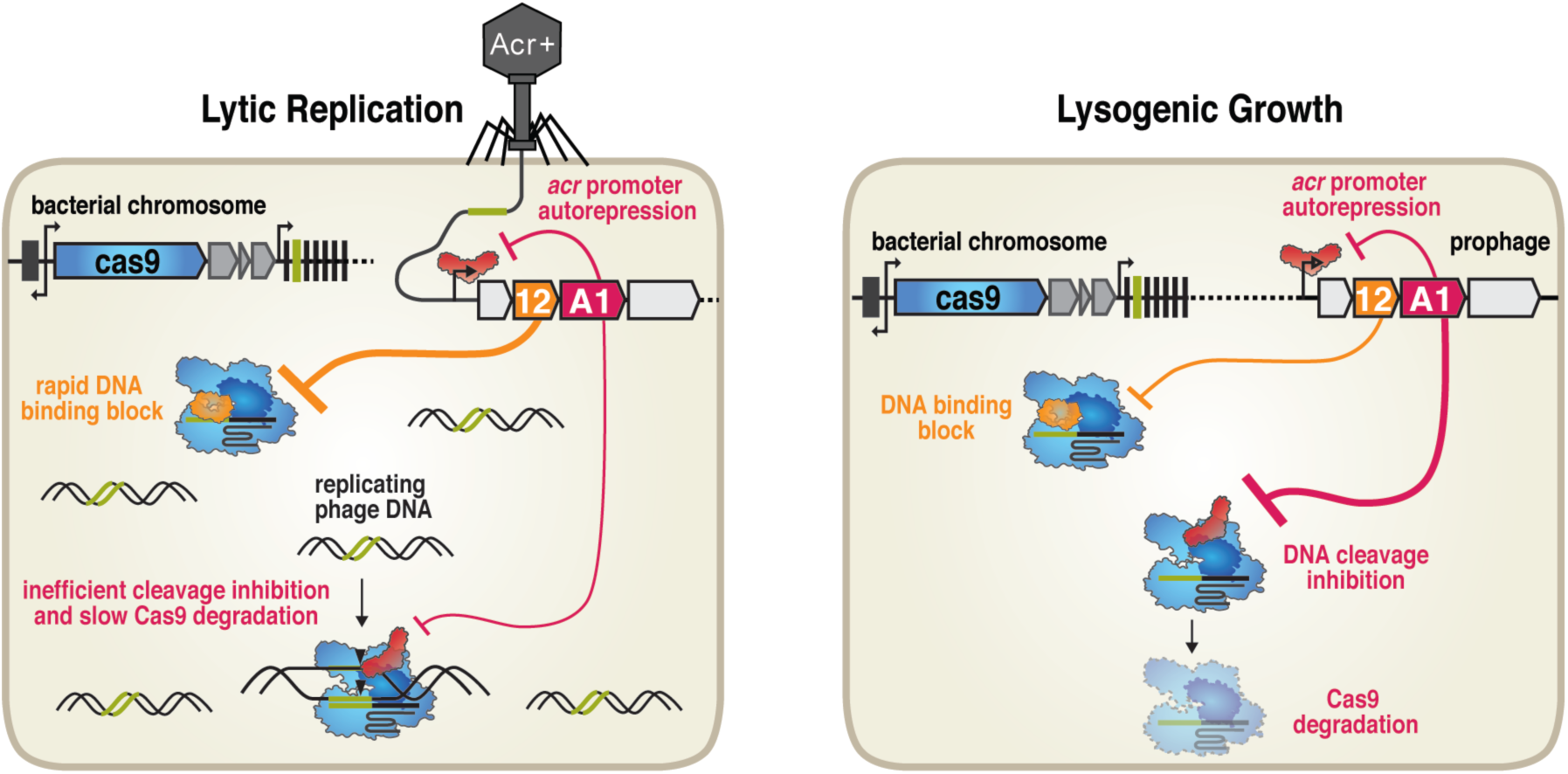
Model for *Listeria* Phage Anti-CRISPR Functions in Lysogenic and Lytic Growth. *Listeria* temperate phages encode the multifunctional AcrIIA1 (red) for protection against CRISPR-Cas in lysogeny (AcrIIA1^CTD^) and autorepression of anti-CRISPR transcription (AcrIIA1^NTD^). In lysogeny (right), AcrIIA1 binds the Cas9 HNH domain (dark blue in Cas9) to prevent DNA cleavage and triggers Cas9 degradation. For replication in lytic growth (left), AcrIIA1 is slow or inefficient, thus a distinct coexisting anti-Cas9 protein (like AcrIIA12, orange) is necessary to rapidly inactivate Cas9.

Notably, this is the first report of an anti-CRISPR that reduces Cas protein levels and is also the first with an additional role integral to the phage life cycle. The highly conserved AcrIIA1^NTD^ plays a general Cas9-independent role by autorepressing *acr* locus transcription to promote phage lytic growth and prophage induction. Engineered phages expressing the AcrIIA1^CTD^ alone had a strong lytic growth defect, perhaps suggesting the AcrIIA1 domains are fused in nature to limit expression of an otherwise problematic anti-CRISPR. Interestingly, when the bacterial host expresses AcrIIA1^NTD^, an “anti-anti-CRISPR” activity manifests, blocking anti-CRISPR expression from infecting or integrated phages. Thus, the importance of the conserved anti-CRISPR locus repression mechanism may represent a weakness that can be exploited by the host through the co-opting of this anti-CRISPR regulator.

Many diverse Cas9 orthologues have been identified and AcrIIA1 can inhibit highly distinct II-A and II-C subtypes. This provides a unique advantage to *Listeria* phages, inhibiting a small LivCas9 variant (25% amino acid identity to large LmoCas9) that is also found in *L. monocytogenes* strains. LivCas9 also shares similarity with Type II-C Cas9s, likely explaining the biological basis of AcrIIA1 activity against the II-C subtypes. Broad-spectrum inhibition by AcrIIA1 is likely due to targeting the highly conserved Cas9 HNH domain catalytic site, whereas the DNA binding inhibitors (AcrIIA2, AcrIIA4, AcrIIA12) are far more limited. AcrIIC1 was similarly reported to block various Type II-C orthologues by directly binding Cas9 (Apo or gRNA-bound) via the HNH domain (Harrington et al., 2017). Much like AcrIIA1, AcrIIC1 binds the NmeCas9 HNH domain with strong affinity (K_D_ = 6.3 nM; Harrington et al., 2017), but it is a rather weak anti-CRISPR in comparison to the DNA binding inhibitors AcrIIC3-5, which have narrow inhibitory spectrums (Lee et al., 2018; Mathony et al., 2019). Therefore, although Cas9 DNA cleavage inhibitors may tend to be weaker anti-CRISPRs, they considerably bolster the phage defense arsenal by targeting a highly conserved, and potentially immutable feature amongst bacterial Cas nucleases. Future engineering of AcrIIA1 could generate a more potent inhibitor, as recently achieved with AcrIIC1 (Mathony et al., 2019). Our attempt to increase anti-CRISPR function in human cells by weakening DNA interactions (AcrIIA1(T16A) mutant, Figure 3D) was only modestly successful.

Widespread AcrIIA1^NTD^ conservation also raises the possibility that prophages use this domain to combat phage superinfection, benefitting both the prophage and host cell. Precedent for phage repressors acting in this manner, both in *cis* and in *trans*, is strong. For example, the phage lambda cI protein represses prophage lytic genes and prevents superinfection by related phages during lysogeny (Johnson et al., 1981). Similarly, lysogens could use AcrIIA1 to temper expression of the prophage anti-CRISPR locus while bolstering the activity of a second CRISPR-Cas system (e.g. Type I-B, which is common in *Listeria*), by preventing incoming phages from expressing their anti-CRISPRs. Given the diversity of anti-CRISPR protein sequences, blocking transcription would be a much more effective strategy than inhibiting individual anti-CRISPRs. Lastly, the widespread nature of the AcrIIA1^NTD^, its fusion to distinct CTDs, and its shared genetic neighborhood with mechanistically distinct anti-CRISPRs, may be a useful marker for future *acr* discovery.

## AUTHOR CONTRIBUTIONS

B.A.O. and J.B.-D. conceived and designed the study. B.A.O., S.Ka., C.M., K.A.C., B.G., and S.Ki. performed experiments. A.R.D., B.P.K., S.Ki., and J.B.-D. supervised experiments. All authors evaluated results. B.A.O. and J.B.-D. wrote the manuscript with input from all authors.

## ACKNOWLEDGEMENTS

We would like to thank Daniel A. Portnoy (UC Berkeley) for providing the pLMB3C-pRhamnose plasmid, Jennifer A. Doudna for Cas9 expression plasmids (UC Berkeley), and Jonathan Asfaha (David Morgan Lab, UCSF) and Ujjwal Rathore (Alex Marson Lab, UCSF) for experimental advice and reagents. The J.B.-D lab was supported by the UCSF Program for Breakthrough Biomedical Research funded in part by the Sandler Foundation, the Searle Fellowship, the Vallee Foundation, an NIH Director’s Early Independence Award DP5-OD021344, and R01GM127489; S.Ki. by an Ambizione Fellowship (Swiss National Science Foundation, SNF_174108); the B.P.K. lab by NIH R00-CA218870 and P01-HL142494, an A.S.G.C.T. Career Development Award, and the Margaret Q. Landenberger Research Foundation; and the A.R.D. lab by a CIHR Foundation grant FDN-15427.

## COMPETING INTERESTS

J.B.-D. is a scientific advisory board member of SNIPR Biome and Excision Biotherapeutics and a scientific advisory board member and co-founder of Acrigen Biosciences. B.P.K. is an inventor on various patents and patent applications that describe gene editing and epigenetic editing technologies, and consults for Avectas Inc.

---

**Figure S1.**
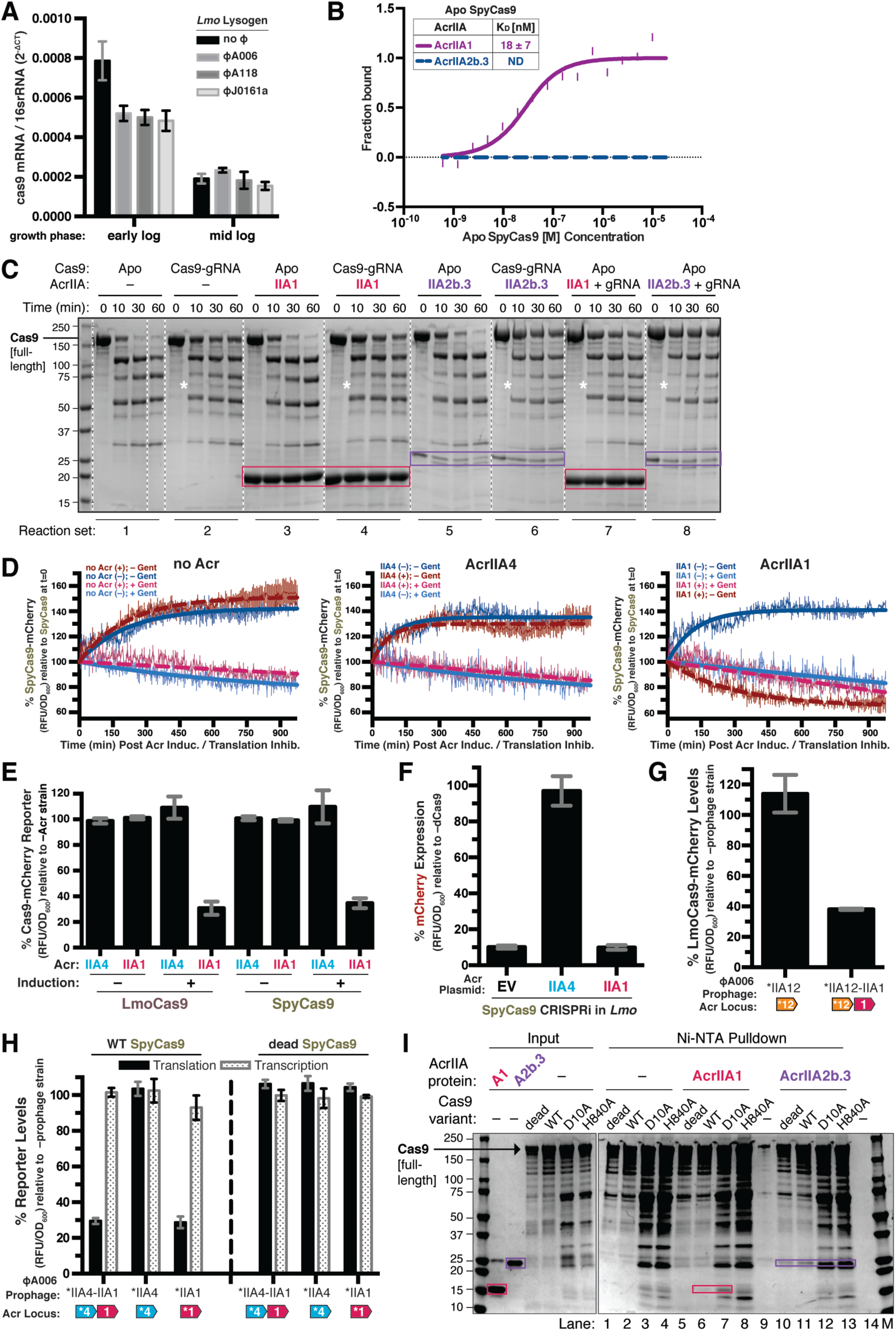
AcrIIA1 Binds Cas9 and Stimulates Post-transcriptional Degradation of Lmo and Spy Cas9 in *Listeria*, Related to Figure 1. (A) Cas9 mRNA levels of *Lmo*10403s lysogens containing the indicated prophages during early or mid-logarithmic growth as quantified by qRT-PCR. Transcript measurements were conducted in technical triplicate and data are shown as the mean 2^−ΔCT^ values normalized to the 16S rRNA endogenous control gene ± SD (error bars). (B) Quantification of the binding affinities (K_D_; boxed inset) of AcrIIA1 and AcrIIA2b.3 for Apo SpyCas9 using microscale thermophoresis. ND indicates no binding was detected. Data shown are representative of three independent experiments. (C) Limited α-chymotrypsin proteolysis of SpyCas9-Acr complexes. Proteolysis of Apo SpyCas9 (set 1) or SpyCas9-gRNA (set 2) without anti-CRISPR (–) or in the presence of AcrIIA1 (sets 3, 4, 7; magenta boxes) or AcrIIA2b.3 (sets 5, 6, 8; purple boxes). For reaction sets 7 and 8, Apo Cas9 was first incubated with anti-CRISPR followed by addition of gRNA. (*) Denotes a proteolysis product that appears in all Cas9-gRNA reactions but not Apo Cas9 reactions. Dashed lines indicate where intervening lanes were removed for clarity. (D) SpyCas9-mCherry protein levels post anti-CRISPR induction or translation inhibition. *Lmo*10403s strains expressing SpyCas9-mCherry from the constitutively active pHyper promoter and AcrIIA1 or AcrIIA4 from an inducible promoter were grown to mid-logarithmic phase and treated with 100 mM rhamnose to induce Acr expression (+, thick dashed lines) or 100 mM glycerol as a neutral carbon source control (–, thick solid lines) and 5 µg/mL gentamicin (Gent) to inhibit translation (+) or water (–) as a control. SpyCas9-mCherry protein measurements reflect the mean percentage fluorescence (RFUs normalized to OD_600_) relative to the SpyCas9-mCherry levels at the time (0 min) translation inhibition was initiated (thin solid lines). Error bars (vertical lines) represent the mean ± SD of at least three biological replicates. Data were fitted by nonlinear regression to generate best-fit decay curves (thick lines). (E) Lmo or Spy Cas9-mCherry protein levels (black bars) in *Lmo*10403s expressing Lmo or Spy Cas9-mCherry from the constitutively active pHyper promoter and AcrIIA1 or AcrIIA4 from an inducible promoter. Cas9-mCherry measurements reflect the mean percentage mCherry (RFU normalized to OD_600_) in cells treated with 100 mM rhamnose (+, induced Acr expression) or 100 mM glycerol (–, non-induced Acr expression), relative to a control strain lacking an anti-CRISPR (–Acr). Error bars represent the mean ± SD of at least three biological replicates. (F) Anti-CRISPR inhibition of CRISPRi in a *Listeria Lmo*10403s strain containing a chromosomally-integrated construct expressing dead SpyCas9 from the inducible pRha-promoter and a sgRNA that targets the pHelp-promoter driving mCherry expression. mCherry expression measurements reflect the mean percentage fluorescence (RFU normalized to OD_600_) in deadCas9-induced cells relative to uninduced (–dCas9) controls of three biological replicates ± SD (error bars). (G) Catalytically active LmoCas9-mCherry protein levels in *Lmo*10403s lysogenized with isogenic ΦA006 prophages encoding AcrIIA12 alone or in combination with AcrIIA1. LmoCas9-mCherry (black bars) measurements reflect the mean percentage mCherry (RFUs normalized to OD_600_) in the indicated lysogens relative to the control strain lacking a prophage (–prophage). Error bars represent the mean ± SD of at least three biological replicates. (*) Indicates the native orfA RBS (strong) in ΦA006 was used for Acr expression. (H) Translational (black bars) and transcriptional (gray shaded bars) reporter levels of catalytically active and dead SpyCas9 in *Lmo*10403s lysogenized with isogenic ΦA006 prophages encoding the indicated anti-CRISPRs. Measurements were normalized and graphed as in (G) with error bars representing the mean ± SD of at least three biological replicates. (*) Indicates the native orfA RBS (strong) in ΦA006 was used for Acr expression. (I) Differential interactions of SpyCas9 nickases with AcrIIA1. Partially purified 6xHis-tagged SpyCas9 (WT, dead, D10A, H840A) proteins (input) were incubated with 2-fold molar excess gRNA and subjected to Ni-NTA pull-down in the presence or absence (lanes 1-4; –) 6-fold molar excess AcrIIA1 (lanes 5-8) or AcrIIA2b.3 (lanes 10-13). AcrIIA1 (magenta boxes) co-purifies with WT and D10A (lanes 6 and 7) but not dead and H840A Cas9-gRNA (lanes 5 and 8). AcrIIA2b.3 (purple boxes) co-purifies with all four Cas9-gRNA complexes (lanes 10-13). AcrIIA1 and A2b.3 were incubated with Ni-NTA beads in the absence of Cas9-gRNA to test for non-specific binding to Ni-NTA beads (lanes 9 and 14).

**Figure S2.**
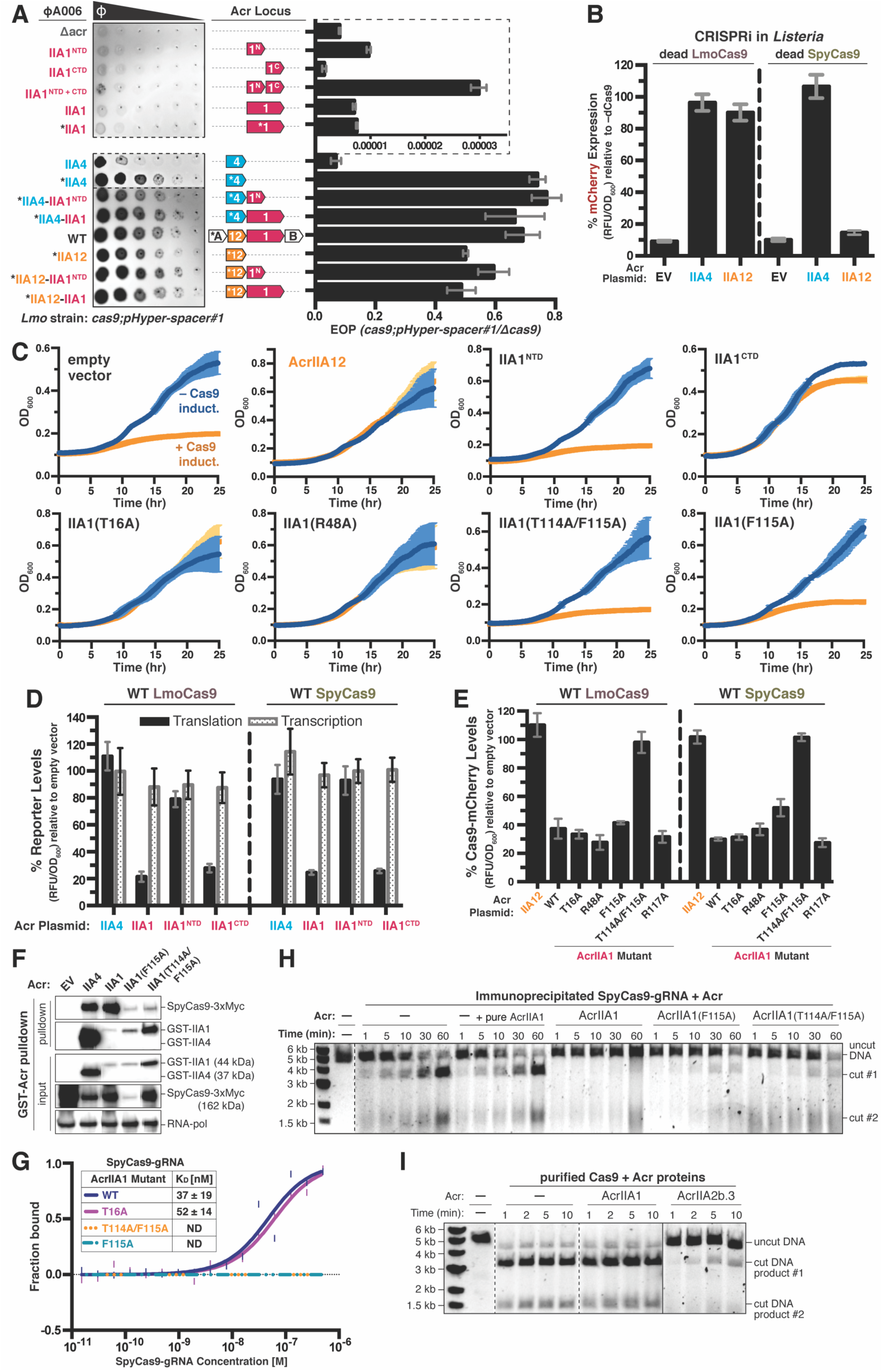
AcrIIA1^CTD^ Mutants Cannot Strongly Bind Cas9 or Trigger its Degradation, Related to Figure 2. (A) Left: Representative image of plaquing assays where isogenic ΦA006 phages are titrated in ten-fold serial dilutions (black spots) on a lawn of *Lmo*10403s (gray background). Dashed lines indicate where intervening rows were removed for clarity. Right: Efficiency of plaquing (EOP) of isogenic ΦA006 phages (expressing the indicated anti-CRISPRs) on *Lmo*10403s. Plaque forming units (PFUs) were quantified on *Lmo*10403s overexpressing the first spacer in the native CRISPR array that targets ΦA006 (*cas9;pHyper-spacer#1*) and normalized to the number of PFUs measured on a non-targeting *Lmo*10403s-derived strain (Δ*cas9*). The dashed lines boxing the first 6 phages show a zoomed in view of the graph with a distinct x-axis scale. Data are displayed as the mean EOP of at least three biological replicates ± SD (error bars). Note that this figure contains the same subset of data displayed in Figure 2A. (B) AcrIIA12 anti-CRISPR activity in a *Lmo*10403s CRISPRi strain expressing Lmo or Spy deadCas9 from the inducible pRha-promoter and a sgRNA that targets the pHelp-promoter driving mCherry expression. mCherry expression measurements reflect the mean percentage RFU in deadCas9-induced cells relative to uninduced (–dCas9) controls of three biological replicates ± SD (error bars). Note that AcrIIA12 inhibits Lmo but not Spy deadCas9-based CRISPRi, indicating its specificity against LmoCas9. (C) Anti-CRISPR activity in *Lmo*10403s self-targeting strains containing chromosomally-integrated constructs expressing LmoCas9 from the inducible pRha-promoter and a sgRNA that targets the pHelp promoter driving mCherry expression. Bacterial growth was monitored after LmoCas9 induction (orange lines) or no induction (blue lines) and data are displayed as the mean OD_600_ of at least three biological replicates ± SD (error bars). (D) Translational (black bars) and transcriptional (gray shaded bars) reporter levels of catalytically active Lmo and Spy Cas9 in *Lmo*10403s containing plasmids expressing anti-CRISPRs. Reporter measurements reflect the mean percentage mCherry (RFU normalized to OD_600_) in the presence of the indicated anti-CRISPRs relative to the control strain containing an empty vector of three biological replicates ± SD (error bars). (E) Catalytically active Lmo and Spy Cas9-mCherry protein levels in *Lmo*10403s containing plasmids expressing anti-CRISPRs. Cas9-mCherry measurements (black bars) reflect the mean percentage mCherry (RFU normalized to OD_600_) in the presence of the indicated anti-CRISPRs relative to the control strain containing an empty vector of three biological replicates ± SD (error bars). (F) Immunoblots detecting 3xMyc-tagged SpyCas9 protein that co-immunoprecipitated with GST-tagged anti-CRISPR proteins in a *P. aeruginosa* strain heterologously expressing the Type II-A SpyCas9-gRNA system and the indicated Acrs. For input samples, one-hundredth lysate volume was analyzed to verify tagged protein expression and RNA-polymerase was used as a loading control. Representative blots of at least three biological replicates are shown. (G) Quantification of the binding affinities (K_D_; boxed inset) of WT and mutant AcrIIA1 proteins with SpyCas9-gRNA using microscale thermophoresis. ND indicates no binding detected. Data shown are representative of three independent experiments. (H-I) Time courses of SpyCas9 DNA cleavage reactions in the presence of the indicated anti-CRISPR proteins conducted with SpyCas9-gRNA-Acr complexes immunoprecipitated from *P. aeruginosa* (H) or recombinant proteins purified from *E. coli* (I). Where indicated, the reaction with SpyCas9-gRNA immunoprecipitated without an Acr (–) was supplemented with recombinant WT AcrIIA1 protein purified from *E. coli* (+ pure AcrIIA1) (H). Dashed lines indicate where intervening lanes were removed for clarity. Solid lines indicate a separate image. Data shown are representative of three independent experiments.

**Figure S3.**
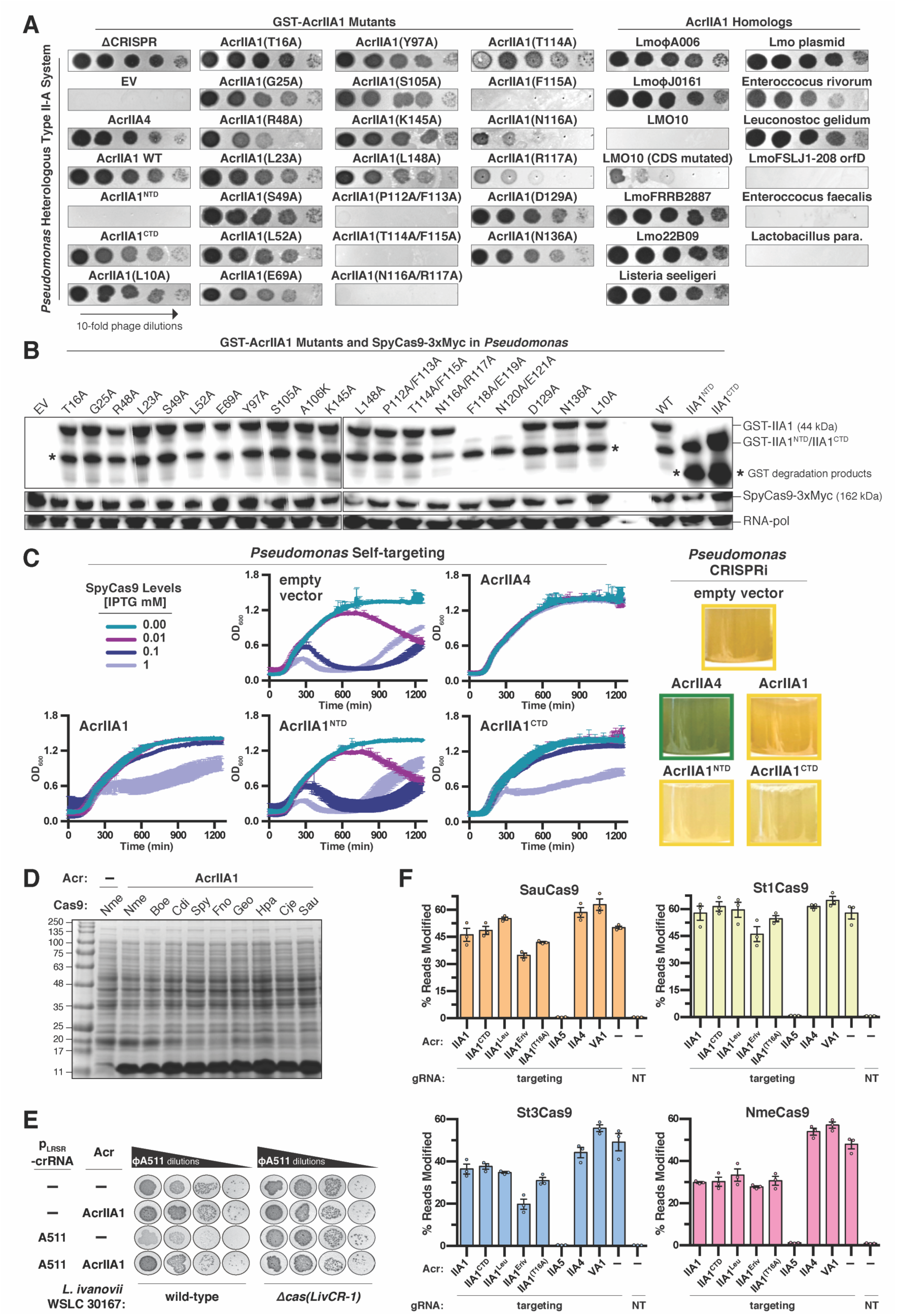
AcrIIA1 Inhibition of Cas9 Orthologues in Heterologous Hosts, Related to Figures 2 and 3. (A) Plaquing assays where the *P. aeruginosa* DMS3m-like phage is titrated in ten-fold dilutions (black spots) on a lawn of *P. aeruginosa* (gray background) expressing the indicated anti-CRISPR proteins and Type II-A SpyCas9-sgRNA programmed to target phage DNA. Representative pictures of at least 3 biological replicates are shown. (B) Immunoblots detecting GST-tagged AcrIIA1 (mutants or individual domains) proteins, Myc-tagged SpyCas9 protein, and RNA-polymerase as a protein loading control in a *P. aeruginosa* strain heterologously expressing the Type II-A SpyCas9-gRNA system and the indicated Acrs. (*) Denotes GST degradation products derived from GST-tagged Acr proteins. AcrIIA1 mutants that failed to express were not analyzed further. (C) Anti-CRISPR activity in *P. aeruginosa* self-targeting (left) and CRISPRi (right) strains containing plasmids expressing anti-CRISPRs and chromosomally-integrated SpyCas9-sgRNA programmed to target the *phZM* gene promoter. For self targeting, SpyCas9 expression from the inducible pLAC-promoter was titrated using the indicated IPTG concentrations (mM) and bacterial growth curves display the mean OD_600_ of at least three biological replicates ± SD (error bars) measured over time (left). CRISPRi was qualitatively assessed by inspecting the culture pigment. Transcriptional repression of the *phzM* gene by dCas9 generates a yellow culture whereas inhibition of dCas9 (e.g. by an Acr) allows *phzM* expression and pyocyanin production that generates a green culture. Representative pictures of at least three biological replicates are shown (right). (D) SDS-PAGE and Coomassie Blue staining analysis of AcrIIA1 expression after IPTG induction in *E. coli* strains containing the indicated Cas9 orthologues. (E) Plaquing assays where the *Listeria* phage ΦA511 is titrated in ten-fold serial dilutions (black spots) on lawns of the *Listeria ivanovii* WSLC 30167 (gray background) strain with an endogenous Type II-A LivCas9 system or lacking this system (Δ*cas*), plasmid-expressed AcrIIA1 or no Acr (–), and crRNA that targets ΦA511 phage DNA or a non-targeting control (–) expressed from the pLRSR plasmid. (F) Gene editing activities of Cas9 orthologues in human cells in the presence of AcrIIA1 variants and orthologues. AcrIIA4 is a known selective inhibitor of SpyCas9, AcrIIA5 is a broad-spectrum inhibitor (Garcia et al., 2019, *in revision*), and AcrVA1 as a known non-inhibiting control for SpyCas9 orthologues. NT, no-sgRNA control condition. Error bars indicate SEM for three independent biological replicates.

**Figure S4.**
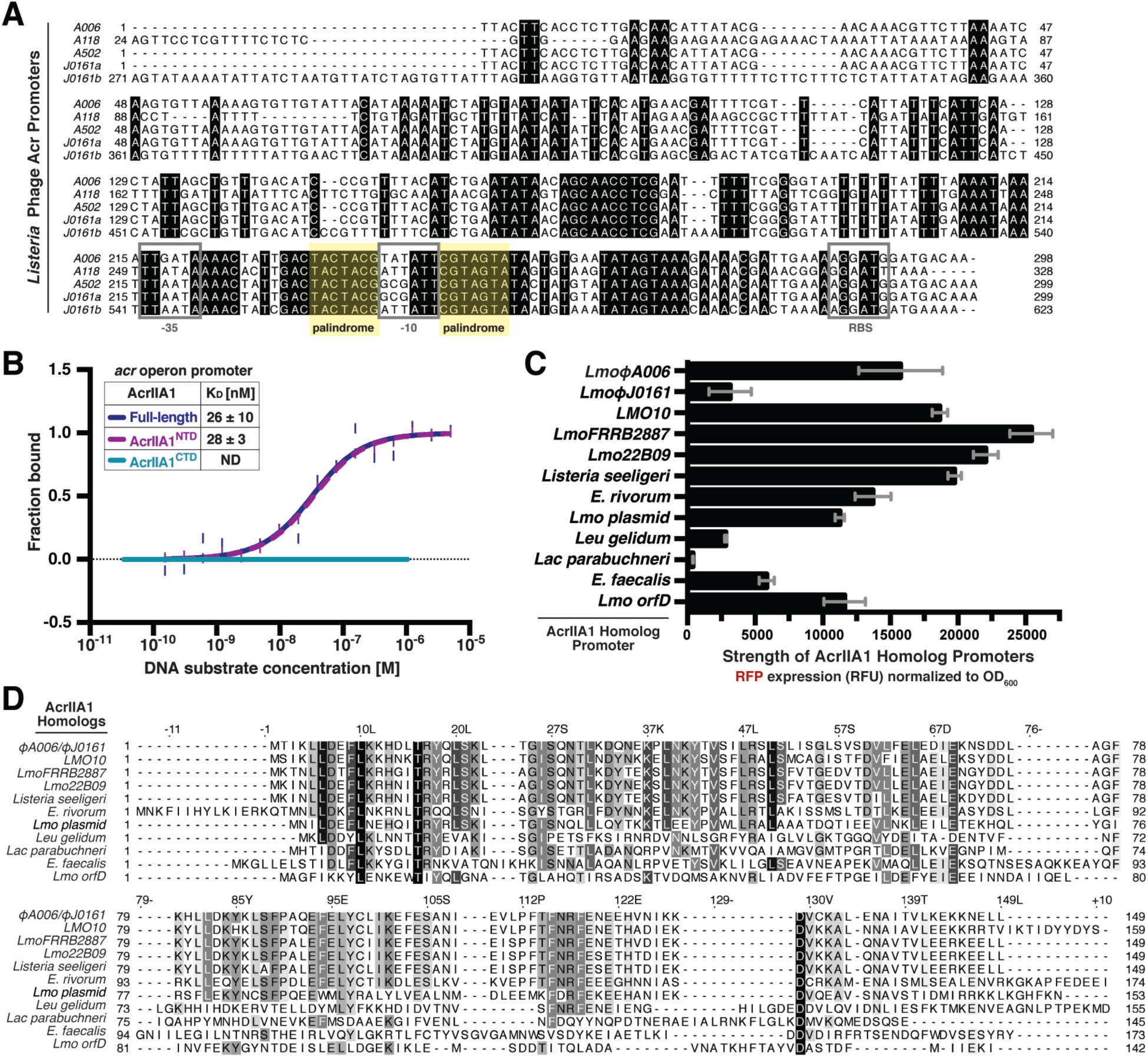
Acr Promoters in Mobile Genetic Elements Across the *Firmicutes* Phylum are Autoregulated by AcrIIA1 Homologs, Related to Figures 2 and 4. (A) Alignment of the phage anti-CRISPR promoter nucleotide sequences denoting the −35 and −10 elements and ribosomal binding site (RBS) (gray boxes) and conserved palindromic sequence (yellow highlight). (B) Quantification of DNA binding abilities (K_D_; boxed inset) of full-length AcrIIA1 and each domain (AcrIIA1^NTD^ and AcrIIA1^CTD^) using microscale thermophoresis. Data shown are representative of three independent experiments. ND indicates no binding detected. (C) Expression strength of the AcrIIA1 homolog promoters. Data are shown as the mean RFP expression (RFU normalized to OD_600_) driven by each AcrIIA1 homolog promoter of at least three biological replicates ± SD (error bars). (D) Alignment of AcrIIA1 homolog protein sequences.

**Figure S5.**
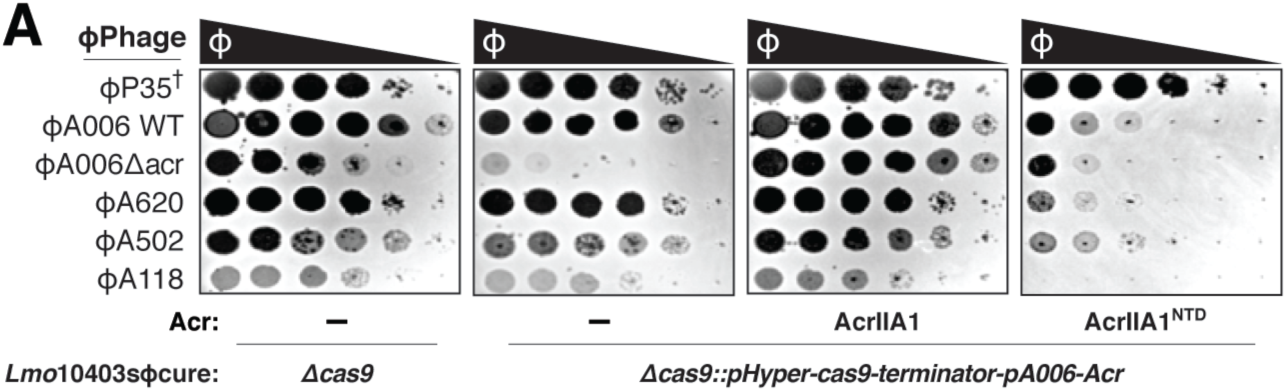
Bacterial expression of AcrIIA1^NTD^ blocks phage anti-CRISPR deployment, Related to Figure 5. (A) Plaquing assays where wild-type *L. monocytogenes* phages are titrated in ten-fold dilutions (black spots) on lawns of *L. monocytogenes* (gray background) containing single-copy integrated constructs expressing AcrIIA1 or AcrIIA1^NTD^ from the ΦA006 anti-CRISPR promoter (pA006), LmoCas9 from a constitutive promoter (pHyper-Cas9), and the natural CRISPR array containing spacers with complete or partial matches to the DNA of each phage. (†) Denotes the absence of a spacer targeting the virulent phage ΦP35. Representative pictures of at least 3 biological replicates are shown.

**Table S1.**
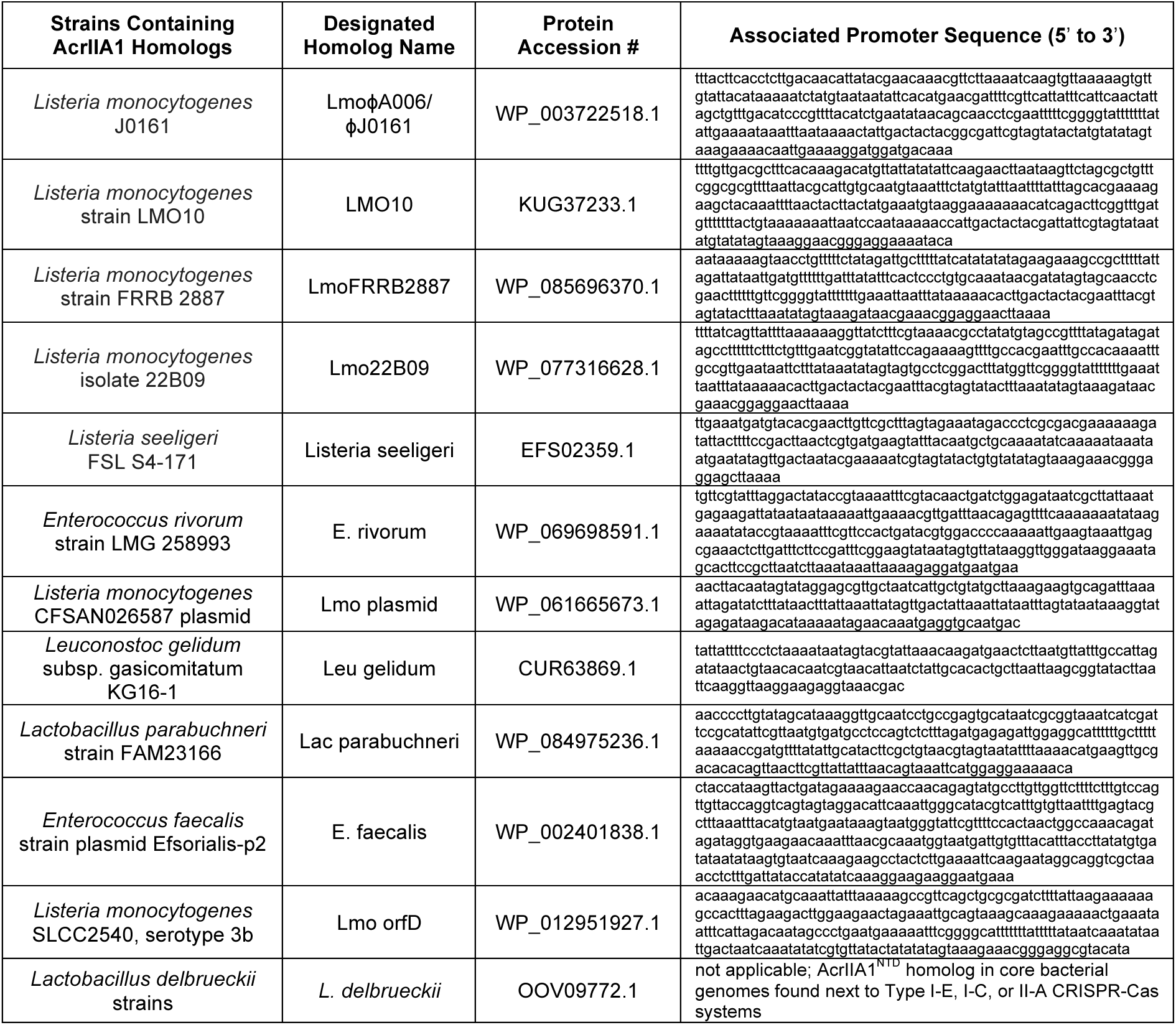
AcrIIA1 homolog protein accession numbers and associated promoter sequences, Related to Figures 2, 4 and 5.

## METHODS

### CONTACT FOR REAGENT AND RESOURCE SHARING

Please direct any requests for further information or reagents to the lead contact, Joseph Bondy-Denomy.

### EXPERIMENTAL MODEL AND SUBJECT DETAILS

#### Microbes

*Listeria monocytogenes* strains (10403s) were cultured in brain-heart infusion (BHI) medium at 30°C. All *Lmo* strains containing pPL2oexL-Rhamnose-inducible constructs were cultured in Luria broth (LB) supplemented with 50-150 mM glycerol (neutral carbon source; no induction/repression) and 0-100 mM rhamnose (inducer) as indicated. To ensure plasmid maintenance in *Listeria* strains, BHI or LB was supplemented with tetracycline (2 µg/mL) for the pPL2oexL integrated construct or erythromycin (7.5 µg/mL) for pLEB579. *Escherichia coli* (DH5α, XL1Blue, NEB 10-beta, or NEB Turbo for plasmid maintenance and SM10 for conjugation into *Listeria*) and *Pseudomonas aeruginosa* (PAO1) were cultured in LB medium at 37°C. To maintain plasmids, LB was supplemented with chloramphenicol (25 µg/mL) for pPL2oexL in *E. coli*, erythromycin (250 µg/mL) for pLEB579 in *E. coli*, gentamicin (30 µg/mL) for pHERD30T in *E. coli* and *P. aeruginosa*, or carbenicillin (250 µg/mL for *P. aeruginosa*, 100 µg/mL for *E. coli*) for pMMB67HE. For maintaining pHERD30T and pMMB67HE in the same *P. aeruginosa* strain, media was supplemented with 30 µg/mL gentamicin and 100 µg/mL carbenicillin.

#### Phages

*Listeria* phages A006, A118, A502, A620, J0161a, and their derivatives were all propagated at 30°C on *acrIIA1^NTD^*-expressing *L. monocytogenes* 10403sɸcure (Δ*cas9*, Δ*tRNAArg::pPL2oexL-acrIIA1^NTD^*) to allow optimal lytic growth of phages lacking their own *acrIIA1^NTD^*. A511 was propagated on *L. ivanovii* WSLC 3009 at 30°C and P35 on *L. monocytogenes* Mack at 20°C. The *Pseudomonas* DMS3m-like phage (JBD30) was propagated on PAO1 at 37°C. All phages were stored in SM buffer (100 mM NaCl, 8 mM MgSO_4_•7H_2_O, 50 mM Tris-HCl pH 7.5, 0.01% (w/v) gelatin), supplemented with 10 mM CaCl_2_ for *Listeria* phages, at 4°C.

#### Human cell lines

Human HEK 293T cells (ATCC) were cultured in Dulbecco’s Modified Eagle Medium (DMEM) supplemented with 10% heat-inactivated FBS (HI-FBS) and 1% penicillin/streptomycin. Media supernatant from cell cultures was analyzed monthly for the presence of mycoplasma using MycoAlert PLUS (Lonza).

### METHOD DETAILS

#### Construction of isogenic ϕA006 anti-CRISPR phages

Isogenic ϕA006 phages encoding distinct anti-CRISPRs from the native anti-CRISPR locus were engineered by rebooting genomic bacteriophage DNA in *L. monocytogenes* L-form cells (EGDe strain variant Rev2) as previously described (Kilcher et al., 2018). Denoted *acr* genes (*) contain the strong ribosomal binding site (RBS) naturally associated with the first gene in the natural ϕA006 anti-CRISPR locus (*orfA*) whereas unmarked genes contain the weaker RBS associated with *acrIIA1*.

#### *Listeria* phage titering

A mixture of 150 µl stationary *Listeria* culture and 3 mL molten LC top agar (10 g/L tryptone, 5 g/L yeast extract, 10 g/L glucose, 7.5 g/L NaCl, 10 mM CaCl_2_, 10 mM MgSO_4_, 0.5% agar) was poured onto a BHI plate (1.5% agar) to generate a bacterial lawn, 3 µl of phage ten-fold serial dilutions were spotted on top, and after 24 hr incubation at 30°C, plate images were collected using the Gel Doc EZ Documentation system (BioRad) and Image Lab (BioRad) software.

#### Construction of *Lmo*10403s::ϕA006/ϕA118/ϕJ0161a lysogens

Lysogens were isolated from plaques that emerged after titering phages ϕA006, ϕA118, ϕJ0161a, and their derivatives on a lawn of *Lmo*10403sɸcureΔ*cas9* (see “*Listeria* phage titering”). Lysogeny was confirmed by prophage induction with mitomycin C (0.5 µg/mL) treatment as previously described (Estela et al., 1992) and by PCR amplification and Sanger sequencing of the phage anti-CRISPR locus. All *Lmo*10403s strains containing prophages were lysogenized and verified prior to introducing additional constructs (integrated pPL2oexL or episomal pLEB579).

#### Construction of L. monocytogenes and P. aeruginosa strains

DNA fragments were PCR-amplified from genomic, plasmid, or synthesized DNA and cloned by Gibson Assembly into *Listeria* plasmids: episomal pLEB579 (Beasley et al., 2004) or the pPL2oexL single-copy integrating plasmid derived from pPL2 (Lauer et al., 2002) or *P. aeruginosa* plasmids: pMMB67HE or pHERD30T. To generate all *Listeria* strains, pPL2oexL plasmids were conjugated (Lauer et al., 2002; Simon et al., 1983) and pLEB579 plasmids were electroporated (Hupfeld et al., 2018; Park and Stewart, 1990) into *Lmo*10403s. For all *Pseudomonas* strains, plasmids were electroporated into PAO1 (Choi et al., 2006).

#### *Listeria* protein samples for immunoblotting

Saturated overnight cultures of *Lmo*10403s strains overexpressing FLAG-tagged Cas9 (Δ*cas9, ΔtRNAArg::pPL2oexL-LmoCas9-6xHis-FLAG*) were diluted 1:10 in BHI with appropriate antibiotic selection (see “microbes”), grown to log phase (OD_600_ 0.2-0.6), harvested by centrifugation at 8000 g for 5 min at 4°C, and lysed by bead-beating or lysozyme treatment. For bead-beating: 4 OD_600_ units of each culture were harvested, cell pellets were resuspended in 500 µl ice cold lysis buffer (50 mM Tris-HCl pH 8.0, 650 mM NaCl, 10 mM MgCl2, 10% glycerol, 1x cOmplete mini EDTA-free protease inhibitor cocktail [Roche]), combined with ∼150 µl 0.1 mm glass beads, and vortexed for 1 hr at 4°C. Cell debris was cleared by centrifugation at 21000 g for 5 min at 4°C and supernatant was mixed with one-third volume 4X Laemmli Sample Buffer (Bio-Rad). For lysozyme lysis: 1.6 OD_600_ units were harvested, cell pellets were resuspended in 200 µl of TE buffer supplemented with 2.5 mg/mL lysozyme and 1x cOmplete mini EDTA-free protease inhibitor cocktail (Roche), samples were incubated at 37°C for 30 min, quenched with one-third volume of 4X Laemmli Sample Buffer (Bio-Rad), and boiled for 5 min at 95°C.

#### Immunoblotting

Protein samples were separated by SDS-PAGE using 4-20% Mini-PROTEAN TGX gels (Bio-Rad) and transferred in 1X Tris/Glycine Buffer onto 0.22 micron PVDF membrane (Bio-Rad). Blots were probed with the following antibodies diluted 1:5000 in 1X TBS-T containing 5% nonfat dry milk: rabbit anti-FLAG (Sigma-Aldrich Cat# F7425, RRID:AB_439687), mouse anti-FLAG (Sigma-Aldrich Cat# F1804, RRID:AB_262044), mouse anti-Myc (Cell Signaling Technology Cat# 2276, RRID:AB_331783), rabbit anti-GST (Cell Signaling Technology Cat# 2625, RRID:AB_490796), mouse anti-*E.coli* RNA polymerase β (BioLegend Cat# 663903, RRID:AB_2564524), HRP-conjugated goat anti-Rabbit IgG (Bio-Rad Cat# 170-6515, RRID:AB_11125142), and HRP-conjugated goat anti-mouse IgG (Santa Cruz Biotechnology Cat# sc-2005, RRID:AB_631736). Blots were developed using Clarity ECL Western Blotting Substrate (Bio-Rad) and chemiluminescence was detected on an Azure c600 Imager (Azure Biosystems).

#### Bacterial growth (OD_600_) and fluorescence (RFU) measurements

Saturated overnight cultures were diluted 1:100 in 150 µl BHI or LB media with appropriate antibiotic selection (see “microbes”) in a 96-well special optics microplate (Corning). *Listeria* cells were incubated at 30°C and *Pseudomonas* at 37°C with continuous double-orbital rotation for 16-48 hr in the Synergy H1 Hybrid Multi-Mode Reader (BioTeK) and measurements of OD_600_ and mCherry (excitation 570 nm, emission 610 nm) or RFP (excitation 555 nm, emission 610 nm) relative fluorescence units (RFU) recorded every 5 min with the Gen5 (BioTek) software. For bacterial growth curves, data are displayed as the mean OD_600_ of at least three biological replicates ± SD (error bars) as a function of time (min or hr, as indicated). For Cas9-mCherry or mCherry fluorescence levels, background fluorescence of growth media was subtracted and the resulting RFU values were normalized to OD_600_ for each time point. Data are displayed as the mean normalized fluorescence 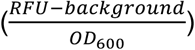 of three biological replicates ± SD.

#### Quantification of Cas9 protein and mRNA reporter levels in *Listeria*

Cas9 (WT or dead; Lmo or Spy) reporters (see Figure 1A schematic) designed to measure protein levels contain a single RBS generating a fused Cas9-mCherry protein. Reporters for mRNA levels contain two ribosomal binding sites, one for Cas9 and a second for mCherry, generating two separate proteins. All reporters were conjugated into *Lmo*10403s devoid of endogenous *cas9* generating strains with the genotype Δ*cas9, ΔtRNAArg::pPL2oexL-pHyper-Cas9Reporter*. Cells were grown and data collected and processed as in “bacterial growth and fluorescence measurements.” Data are shown as the percentage of Cas9 translation and transcription levels (mCherry fluorescence averaged across 6 hr of logarithmic growth) relative to control strains (no prophage (–prophage) or empty vector, as indicated) of at least three biological replicates ± SD (error bars).

#### RT-qPCR of *cas9* mRNA levels

WT or Cas9-overexpressing *Lmo*10403s (Δ*cas9, ΔtRNAArg::pPL2oexL-LmoCas9-6xHis-FLAG*) strains were grown to early (OD_600_ 0.2-0.3) or mid-log (OD_600_ 0.4-0.6) phase and 1.6 OD_600_ units of cells were harvested as in “*Listeria* protein samples.” Cell pellets were resuspended in 100 µl TE buffer supplemented with 0.2 U/µl SUPERase•In RNase Inhibitor (Thermo Fisher Scientific) and 5 mg/mL lysozyme, and incubated at 37°C for 10 min. Each sample was mixed with solutions pre-heated to 65°C for 15 min: 600 µl hot 1.2X lysis buffer (60 mM NaOAc, 1.2% SDS, 12 mM EDTA) and 700 µl hot acid-phenol:chloroform pH 4.5 (with IAA, 125:24:1) (Ambion). After incubating at 65°C for 30 min with shaking at 1500 rpm, followed by centrifugation at 12000 g for 15 min at 4°C, 500 µl aqueous phase was recovered for each sample. RNA was extracted with neutral phenol:chloroform:isoamyl alcohol (25:24:1) (Sigma) three times, precipitated with ethanol, and resuspended in nuclease-free water. Residual DNA was removed using the TURBO DNA-free Kit (Invitrogen). RT-qPCR was conducted in technical triplicate using the Luna Universal One-Step RT-qPCR Kit (New England Biolabs) according to the manufacturer’s instructions in 10 µl reaction volumes and reactions were run on a CFX Real-Time PCR Detection System (BioRad). *cas9* mRNA and *16srRNA* were analyzed with the following primers: cas9-FWD: 5′-ATGCCGCGATAGATGGTTAC-3′ and cas9-REV: 5′-CGCCTTCGATGTTCTCCAATA-3′; 16s-FWD: 5′-CCTGGTAGTCCACGCCGT-3′ and 16s-REV: 5′-TGCGTTAGCTGCAGCACTAAG-3′.

#### Cas9 and anti-CRISPR protein expression and purification

N-terminally 6xHis-tagged Acr proteins were expressed from the pET28 vector whereas WT SpyCas9 and mutants were expressed from 6xHis-MBP-Cas9 constructs (gifts from Jennifer Doudna, UC Berkeley) in *E. coli* Rosetta (DE3) pLysS cells. Recombinant protein expression was induced with 0.25 mM isopropyl β-D-1-thiogalactopyranoside (IPTG) at 18 °C overnight. Cells were harvested by centrifugation and lysed by sonication in buffer A (50 mM Tris-HCl pH 7.5, 500 mM NaCl, 0.5 mM DTT, 20 mM imidazole, 5% glycerol) supplemented with 1 mM PMSF and 0.25 mg/mL lysozyme (Sigma). Cell debris was removed by centrifugation at 20000 g for 40 min at 4 °C and the lysate incubated with Ni-NTA Agarose Beads (Qiagen). After washing, bound proteins were eluted with Buffer A containing 300 mM imidazole and dialyzed overnight into storage buffer (20 mM HEPES-NaOH pH 7.4, 150mM KCl, 10% glycerol, 2mM DTT). GST-tagged AcrIIA1 and AcrIIA2b.3 were expressed from pGEX-6P-1 plasmids in *E. coli* BL21 (DE3) cells, lysed in buffer (20 mM HEPES-NaOH pH 7.4, 300 mM KCl and 5 mM DTT) supplemented with 1 mM PMSF and 0.25 mg/mL lysozyme, and clarified lysate was incubated with Glutathione Agarose Beads (Pierce). After washing, bound proteins were eluted using 100 mM Tris-HCl pH 8.5, 150 mM KCl, 15 mM reduced glutathione. The GST tag was cleaved with PreScission Protease (Millipore) and proteins were dialyzed overnight in 50 mM HEPES-NaOH pH 7.5, 150 mM KCl, 10% glycerol and 2 mM DTT to remove free glutathione. Cleaved GST was removed from dialyzed proteins with Glutathione Agarose Beads (Pierce).

#### *in vitro* binding of anti-CRISPRs to SpyCas9

The binding affinities of anti-CRISPR proteins to SpyCas9 were calculated using microscale thermophoresis (MST) on the Monolith NT.115 instrument (NanoTemper Technologies GmbH, Munich, Germany). For AcrIIA1/AcrIIA2b.3 with WT or mutant Cas9-gRNA complexes, WT and mutant 6xHis-Cas9 proteins were incubated with two-fold molar excess gRNA (Integrated DNA Technologies). The substrate proteins AcrIIA1/AcrIIA2b.3 at 0.09 nM to 3 µM concentrations were incubated with 25 nM RED-tris-NTA-labeled 6xHis-Cas9-gRNA at room temperature (RT) for 10 min in MST buffer (50 mM Tris-HCl pH 7.4, 150 mM NaCl, 15 mM MgCl_2_, 0.05% Tween-20). For AcrIIA1/AcrIIA2b.3 with apoCas9, the substrate protein apoCas9 (QB3 Macrolab) at 0.61 nM to 10 µM concentrations was incubated with 25 nM NT-647-NHS-labeled AcrIIA1/A2b.3 proteins at RT for 10 min in MST buffer. For AcrIIA1 mutants with WT Cas9-gRNA, the substrate protein Cas9-gRNA (QB3 Macrolab) at 15 pM to 0.5 µM concentrations was incubated with 25 nM RED-tris-NTA-labeled 6xHis-AcrIIA1 mutant proteins at RT for 10 min in MST buffer. Samples were loaded into Monolith NT.115 Capillaries and measurements were performed at 25 °C using 40% LED power and medium microscale thermophoresis power. All experiments were repeated three times for each measurement. Data analyses were carried out using NanoTemper analysis software.

#### *in vitro* pull-downs to verify binding of anti-CRISPRs to SpyCas9

5 µg apoCas9 proteins (WT, dead, D10A, or H840A) were incubated with two-fold molar excess gRNA at 37°C for 15 min. Cas9-gRNA complexes were incubated with 6-fold molar excess AcrIIA1 or AcrIIA2b.3 proteins for 15 min at room temperature in a buffer containing 20 mM HEPES-NaOH pH 7.5, 150 mM KCl, 10% glycerol, and 1 mM DTT. Samples were then incubated with 20 µl Ni-NTA agarose beads (Qiagen) for 15 min at 4°C and washed five times with 1x MST buffer (50 mM Tris-HCl pH 7.4, 150 mM NaCl, 10 mM MgCl2, 0.05 % Tween-20). Beads were boiled in 1X Laemmli Sample Buffer and proteins were analyzed by SDS-PAGE and Bio-Safe Coomassie staining (Biorad).

#### Limited proteolysis of SpyCas9-AcrIIA1 complex

20 µg purified SpyCas9 (QB3 Macrolab) in Apo form or in complex with gRNA (1.1-fold molar excess) was incubated with 1.5-fold and 4-fold molar excess AcrIIA1 and AcrIIA2b.3, respectively, in protease buffer (10 mM Tris-HCl pH 7.5, 300 mM NaCl) at 25°C for 15 min. Alternatively, ApoSpyCas9 was incubated first with AcrIIA protein followed by gRNA addition. Proteolysis reactions were performed with 20 ng α-chymotrypsin (sequencing grade, Promega) at 25°C and at 0, 10, 30, or 60 min time points, reactions were quenched with 2X SDS Laemmli Buffer and boiled for 10 min at 95°C. Samples were analyzed by SDS-PAGE and staining with Bio-Safe Coomassie (Bio-Rad).

#### SpyCas9 protein decay measurements in *Listeria*

Saturated overnight cultures of *Lmo*10403s strains devoid of endogenous *cas9* and expressing AcrIIA1 or AcrIIA4 from a tightly regulated rhamnose-inducible promoter (Fieseler et al., 2012) and SpyCas9-mCherry from the constitutively active pHyper promoter (Δ*cas9, ΔtRNAArg::pPL2oexL-pHyper-SpyCas9-mCherry-GyrA_terminator-pRha-AcrIIA*) were diluted 1:100 in fresh LB supplemented with 50 mM glycerol and tetracycline (2 µg /mL) and grown to mid-log (OD_600_ ∼0.5). Cultures were then diluted 1:2 in LB containing 50 mM glycerol and tetracycline (2 µg /mL) plus 200 mM rhamnose to induce Acr expression or 200 mM glycerol for uninduced controls (100 mM final concentration rhamnose or glycerol) in a 96-well microplate and treated with gentamicin (5 µg/mL) to inhibit translation or water as a control. Cells were grown and data collected and processed as in “bacterial growth and fluorescence measurements.” Data are shown as the mean percentage of SpyCas9-mCherry fluorescence relative to levels measured at “0 hr” (the beginning of translation inhibition or anti-CRISPR induction) of at least three biological replicates ± SD (error bars) as a function of time (min). Data were fitted by nonlinear regression to generate best-fit decay curves.

#### *Listeria* CRISPRi and self-targeting

Single-copy integrating CRISPRi and self-targeting constructs (see Figure 1E schematics) were designed as follows: pPL2oexL–pHyper-sgRNA [pHELP-spacer] GyrATerminator–pRhamnose-Cas9 (Lmo WT or Lmo dead or Spy dead) LambdaTerminator–pHELP-mCherry-LuxTerminator and conjugated into *Lmo*10403sɸcureΔ*cas9* containing pLEB579 plasmids expressing the indicated anti-CRISPRs. Overnight cultures were grown in LB supplemented with 50 mM glycerol (no induction/repression), 2 µg/mL tetracycline, and 7.5 µg/mL erythromycin. Cultures were then diluted 1:100 in LB containing 50 mM glycerol and the aforementioned antibiotics plus 200 mM rhamnose to induce Cas9 expression (and thus, CRISPRi or self-targeting) or 200 mM glycerol for uninduced controls (100 mM final concentration rhamnose or glycerol) in a 96-well microplate. Cells were grown and data collected and processed as in “bacterial growth and fluorescence measurements.” For self-targeting, data are displayed as the mean OD_600_ of at least three biological replicates ± SD (error bars) as a function of time (hr). For CRISPRi, data are shown as the mean percentage mCherry expression (mCherry fluorescence averaged across 6 hr of logarithmic growth) relative to uninduced controls of at least three biological replicates ± SD (error bars).

#### Plaque forming unit (PFU) quantification of *Listeria* phages

Phage infections were conducted using the soft agar overlay method: 10 µl phage dilution was mixed with 150 µl stationary *Listeria* culture in 3 mL molten LC top agar supplemented with 300 µg/mL Tetrazolium Violet (TCI Chemicals) to generate contrast for plaque visualization (Hurst et al., 1994) and poured onto a BHI-agar plate. After 24 hr incubation at 30°C, phage plaque-forming units (PFU) were quantified.

#### Efficiency of plaquing of *Listeria* phages

Efficiency of plaquing (EOP) calculations are a ratio of the number of plaque forming units (PFUs) that formed on a *Lmo*10403sɸcure targeting strain (endogenous *cas9* with overexpression of the native CRISPR array spacer #1 that targets ɸA006) divided by the number of PFUs that formed on a non-targeting strain (Δ*cas9*). Each PFU measurement was conducted in biological triplicate and all EOP data is displayed as the mean EOP ± SD (error bars).

#### Construction of self-targeting 10403s::ϕA006 lysogens

*Lmo*10403sΔ*cas9*::ϕA006 isogenic self-targeting lysogens encoding no anti-CRISPR or AcrIIA1, AcrIIA4, AcrIIA12 (alone or in combination as indicated) were isolated as in “construction of *Lmo*10403s lysogens.” To prevent self-targeting during strain construction, pPL2oexL constructs encoding a tightly regulated rhamnose-inducible LmoCas9 (WT or dead as a control) were conjugated into each lysogen. To assess the stability of each lysogen, cells were cultured, Cas9 induced, and data displayed as described for the self-targeting strain in “*Listeria* CRISPRi and self-targeting,” except erythromycin was omitted from LB media. Each lysogen stability measurement was performed in biological triplicate.

#### *P. aeruginosa* anti-SpyCas9 screening platform

The previously described *P. aeruginosa* anti-SpyCas9 screening platform (Jiang et al., 2019) and bacteriophage plaque assays (Borges et al., 2018; Jiang et al., 2019) were utilized to assay the anti-CRISPR activity of AcrIIA1 homologs and mutants. AcrIIA1 homolog genes were synthesized (Twist Bioscience) and cloned into the pMMB67HE-P_Lac_ vector. Protein accession numbers are listed in Table S1. Site directed mutagenesis by Gibson Assembly was used to introduce point mutations into pMMB67HE-P_Lac_-GST-AcrIIA1. The P_BAD_ promoter driving chromosomally integrated SpyCas9-3xMyc and pHERD30T-sgRNA was induced with 0.1% arabinose and the P_Lac_ promoter driving pMMB67HE-AcrIIA with 1 mM IPTG. Expression of AcrIIA1 mutants was confirmed by harvesting 1 OD_600_ unit of cells and resuspending in 200 µl 1X Laemmli Sample Buffer (Bio-Rad) followed by SDS-PAGE and immunoblotting as described above. The fold reductions in phage titer displayed were qualitatively derived by examining at least three replicates of each experiment. Plate images were acquired as in “*Listeria* phage titering” and a representative picture is shown.

#### *P. aeruginosa* self-targeting and CRISPRi

Strains were generated as previously described by Borges et al., 2018 under “construction of PAO1::SpyCas9 expression strain,” except the sgRNA was designed to target the PAO1 chromosomal *phzM* gene promoter and was integrated into the bacterial genome using the mini-CTX2 vector (Hoang et al., 2000). Cultures were grown overnight in LB supplemented with 50 µg/mL gentamicin and 0.1% arabinose to pre-induce anti-CRISPR expression and the next day diluted 1:100 with fresh LB containing 50 µg/mL gentamicin, 0.1% arabinose, and IPTG (0, 0.01, 0.1 or 1mM to titrate WT or dead SpyCas9-sgRNA expression) in a 96-well microplate (150 µl/well) for self-targeting analysis or glass tubes (3 mL) for CRISPRi. Self-targeting experiments were conducted in biological triplicate with cells grown and data collected and processed as in “bacterial growth and fluorescence measurements.” For CRISPRi, cells were grown for 8-10 hr with continuous shaking after which CRISPRi was qualitatively assessed by inspecting the culture pigment. Repression of the *phzM* gene by dCas9 generates a yellow culture whereas inhibition of dCas9 (e.g. by an Acr) allows *phzM* expression and pyocyanin production that generates a green culture. Representative pictures of at least three biological replicates are shown.

#### Co-immunoprecipitation of SpyCas9-3xMyc and GST-AcrIIA

Saturated overnight cultures of *P. aeruginosa* strains were diluted 1:100 in 50 mL of LB supplemented with required antibiotics, grown to OD_600_ 0.3-0.4, and induced with 0.3% arabinose (SpyCas9-gRNA) and 1mM IPTG (anti-CRISPR). Cells were harvested at OD_600_ 1.8-2.0 by centrifugation at 6000 g for 10 min at 4°C, flash frozen on dry ice, resuspended in 1 mL lysis buffer (50 mM Tris-Cl pH 7.4, 150 mM NaCl, 20 mM MgCl_2_, 0.5% NP-40, 5% (v/v) glycerol, 5 mM DTT, 1 mM PMSF), lysed by sonication (20 sec pulse × 4 cycles with cooling between cycles), and lysate was clarified by centrifugation at 14 000g for 10 min at 4°C. For input samples, 10 µL lysate was mixed with one-third volume 4X Laemmli Sample Buffer. Remaining lysate (∼1 mL) was mixed with pre-washed Myc-Tag Magnetic Bead Conjugate #5698 (Cell Signaling Technology) or Glutathione Magnetic Agarose Beads #78601 (Thermo Fisher Scientific) using a lysate to bead slurry volume ratio of 20:1 for Myc or 40:1 for GST. After overnight incubation at 4°C with end-over-end rotation, beads were washed five times with 1 mL cold wash buffer (50 mM Tris-HCl pH 7.4, 150 mM NaCl, 20 mM MgCl_2_, 5mM DTT) containing decreasing concentrations of NP-40 (0.5%, 0.05%, 0.01%, 0.005%, 0) and glycerol (5%, 0.5%, 0.05%, 0.005%, 0) on a magnetic stand. Bead-bound proteins were resuspended in 100 µl wash buffer without detergent and glycerol. 10 µl bead-bound protein slurry was mixed with one-third volume 4X Laemmli Sample Buffer, boiled for 5 min at 95°C, and samples were analyzed by SDS-PAGE using 4-20% Mini-PROTEAN TGX gels (Bio-Rad) and staining with Bio-Safe Coomassie (Bio-Rad) or immunoblotting.

#### Cas9 DNA cleavage assays using immunoprecipitated SpyCas9-3xMyc

Reactions were assembled with bead-bound protein slurry and 1.5 nM DNA substrate, incubated at 25°C with gentle shaking at 1000 rpm, and at 1, 5, 10, and 30 min time points reaction aliquots were mixed with warm Quenching Buffer (50 mM EDTA, 0.02% SDS) and boiled at 95°C for 10 min. DNA cleavage products were analyzed by agarose (1%) gel electrophoresis and staining with SYBR Safe (Thermo Fisher Scientific).

#### Cas9 DNA cleavage assays using purified proteins

To generate gRNAs, crRNA and tracrRNA were annealed with Nuclease-free Duplex Buffer (Integrated DNA Technologies) according to the manufacturer’s instructions. Reactions were assembled in 1X MST Buffer (50 mM Tris-Cl pH 7.4, 150 mM NaCl, 20 mM MgCl_2_, 5 mM DTT, 5% glycerol, 0.05% (v/v) Tween-20) with 50 nM SpyCas9 and 625 nM AcrIIA, incubated for 5 min on ice, supplemented with 50 nM gRNA, and incubated for an additional 5 min at room temperature. Reactions were initiated by adding 2 nM target DNA substrate and at 1, 2, 5 and 10 min time points reaction aliquots were mixed with warm Quenching Buffer (50mM EDTA, 0.02% SDS) and boiled at 95°C for 10 min. DNA cleavage products were analyzed by agarose (1%) gel electrophoresis and staining with SYBR Safe (Thermo Fisher Scientific).

#### *E. coli* phage Mu plaquing assays

Plasmids expressing Type II-A, II-B, and II-C Cas9-sgRNA combinations were previously described (Garcia et al., 2019, *in revision*). Cas9 plasmids containing a spacer targeting phage Mu and a pCDF-1b plasmid expressing the indicated anti-CRISPR proteins were co-transformed into *E. coli* BB101. After 2 hr of growth in LB at 37°C with continuous shaking, cells were treated with 0.01 mM IPTG to induce anti-CRISPR expression, and incubated for an additional 3 hr. A mixture of cells and LB top agar (0.7% agar) was poured onto an LB plate supplemented with 200 ng/mL aTc, 0.2% arabinose, and 10 mM MgSO_4_. Ten-fold serial dilutions of phage Mu were spotted on top and plates were incubated overnight. Anti-CRISPR expression after IPTG induction was analyzed by SDS-PAGE on a 15% Tris-Tricine gel followed by Coomassie Blue staining as previously described (Lee et al., 2018).

#### Inhibition of *LivCas9* by anti-CRISPR proteins

Plaquing assays were conducted as previously described by Hupfeld et al., 2018. Briefly, a pKSV7-derived plasmid expressing AcrIIA1 from the ΦA006 anti-CRISPR promoter or empty vector and pLRSR-crRNA plasmids with a spacer against phage ΦP35, ΦA511, or a non-targeting control were transformed into a *Listeria monocytogenes* Mack strain containing chromosomally-integrated pHelp-LivCas9/tracrRNA or a *Listeria ivanovii* WSLC 30167 strain with an endogenous Type II-A LivCas9 system. A mixture of 200 µl stationary host culture and 4 mL LC top agar was poured onto an agar plate (LC for ΦP35; 1/2 BHI for ΦA511). Ten-fold serial dilutions of phage were spotted on top, plates were incubated at 20°C for ΦP35 and 30°C for ΦA511 for one day, and plate images were subsequently acquired.

#### Generation of human cell expression plasmids

Descriptions of plasmids used for expression of sgRNAs (including sgRNA/crRNA target sequences), nucleases, and Acr proteins in human cells are available upon request. U6 promoter sgRNA and crRNA expression plasmids were generated by annealing and ligating oligonucleotide duplexes into BsmBI-digested BPK1520, BPK2660, KAC14, KAC27, KAC482, KAC32 and BPK4449 for SpyCas9, SauCas9, St1Cas9, St3Cas9, CjeCas9, and NmeCas9, respectively. New human cell expression plasmids for CjeCas9, St3Cas9, and NmeCas9 were generated by sub-cloning the nuclease open-reading frames of Addgene plasmids # 89752, 68337, and 119923, respectively (gifts from Seokjoong Kim, Feng Zhang and Erik Sontheimer) into the AgeI and NotI sites of pCAG-CFP (Addgene plasmid 11179; a gift from C. Cepko). Human codon optimized Acr constructs containing a C-terminal SV40 nuclear localization signal were generated by isothermal assembly of synthetic gene fragments (Twist Biosciences) into the NotI and AgeI sites of Addgene plasmid ID 43861. New human expression plasmids described in this study have been deposited with Addgene.

#### Transfection of human cells

Approximately 20 hours prior to transfection, HEK 293T cells were seeded at 2×10^4^ cells/well in 96-well plates. Cells were transfected using 70 ng of nuclease, 30 ng sgRNA/crRNA, and 110 ng of anti-CRISPR expression plasmids with 1.25 µl of *Trans*IT-X2 (Mirus Bio) in 20 µl Opti-MEM. For control conditions containing no acr plasmid, 110 ng of a pCMV-EGFP plasmid was utilized as filler DNA; for non-targeting sgRNA/crRNA conditions, 30 ng of an empty U6 promoter plasmid was used as filler DNA. Genomic DNA was harvested from cells 72 hours post-transfection by suspending cells in 100 µl of lysis buffer (20 mM Hepes pH 7.5, 100 mM KCl, 5 mM MgCl_2_, 5% glycerol, 25 mM DTT, 0.1% Triton X-100, and 30 ng/ul Proteinase K (NEB)), followed by incubation at 65°C for 6 minutes and 98°C for 2 minutes. All experiments were performed with at least 3 independent biological replicates.

#### Assessment of Cas and Acr protein activities in human cells

Genome editing efficiencies were determined by next-generation sequencing using a 2-step PCR-based Illumina library construction method. Briefly, genomic regions were initially amplified using Q5 High-Fidelity DNA Polymerase (NEB), ∼100 ng of genomic DNA lysate, and gene-specific round 1 primers. PCR products were purified using paramagnetic beads as previously described (Kleinstiver et al., 2019) and diluted 1:100 prior to the 2^nd^ round of PCR to add Illumina barcodes and adapter sequences using Q5 polymerase. PCR amplicons were bead purified, quantified and normalized (Qiagen QIAxcel), and pooled. Final libraries were quantified using an Illumina Library qPCR Quantification Kit (KAPA Biosystems) and sequenced on a MiSeq sequencer using a 300-cycle v2 kit (Illumina). Genome editing activities were determined from the sequencing data using CRISPResso2 (Clement et al., 2019) with commands --min_reads_to_use_region 100 and -w 10.

#### Quantification of prophage induction efficiency

Prophages were induced from *Lmo*10403s::ΦJ0161 lysogens expressing *cis-acrIIA1* from the prophage Acr locus or *trans-acrIIA1* from the bacterial host genome by treating with 0.5 µg/mL mitomycin C as previously described (Estela et al., 1992). After overnight incubation with continuous shaking at 30°C, cells were pelleted by centrifugation at 8000 g for 10 min and phage-containing supernatants were harvested. To quantify the amount of phage induced from each lysogen, phage-containing supernatants were used to infect *Lmo*10403sΦcure lacking *cas9* and expressing AcrIIA1^NTD^ (Δ*cas9;IIA1^NTD^*, to bypass the lytic growth defect of ΦJ0161Δ*acrIIA1-2*) as described in “plaque forming unit (PFU) quantification of *Listeria* phages” and the resulting PFUs were quantified. Data are displayed as the mean PFU/mL after prophage induction of four biological replicates ± SD (error bars).

#### Transcriptional repression of the *acr* promoter

To generate *acr* promoter transcriptional reporters, the nucleotide sequences (∼100-350 base pairs) upstream of putative *acr* loci encoding *acrIIA1* homologs were synthesized (Twist Bioscience) and cloned upstream of an mRFP gene into the pHERD30T vector. Promoter sequences are listed in Table S1. Transcriptional reporters were electroporated into *P. aeruginosa* PAO1 strains containing pMMB67HE-AcrIIA1-variants. Saturated overnight cultures were diluted 1:10 in LB supplemented with 30 µg/mL gentamicin, 100 µg/mL carbenicillin, and 1 mM IPTG to induce AcrIIA1 expression in a 96-well microplate. Cells were grown and data collected as in “bacterial growth and fluorescence measurements.” Data are shown as the mean percentage RFP repression (RFU values at 960 min for AcrIIA1 mutants and 1170 min for homologs, normalized to OD_600_) in the presence of AcrIIA1 relative to controls lacking AcrIIA1 of at least three biological replicates ± SD (error bars).

#### *in vitro* binding of AcrIIA1 to the anti-CRISPR promoter

The affinities of AcrIIA1 and individual domains for DNA were measured in triplicate using MST as described above. Single-stranded complementary oligonucleotides were annealed to generate 40 bp *acr* promoter fragments harboring WT or mutated palindrome. The DNA substrate at 0.15 nM to 5 µM concentrations was incubated with 12.5 nM RED-tris-NTA-labeled AcrIIA1/domains at room temperature for 10 min in 1x buffer (50 mM Tris-HCl pH 7.4, 150 mM NaCl, 10 mM MgCl2, 0.05 % Tween-20). DNA substrate sequences used are as follows:

5’-AACTATTGAC**TACTACG**TATATT**CGTAGTA**TAATGTGAAT-3’ (Wild-type)

5’-AACTATTGAC**<u>A</u>ACTACG**TATATT**CGTAGT<u>T</u>**TAATGTGAAT-3’ (Terminal Mutations)

5’-AACTATTGAC**<u>A</u>AC<u>A</u>AC<u>C</u>**TATATT**<u>G</u>GT<u>T</u>GT<u>T</u>**TAATGTGAAT-3’ (Six Mutations)

#### QUANTIFICATION AND STATISTICAL ANALYSIS

All numerical data, with the exception of the microscale thermophoresis (MST) data, were analyzed and plotted using GraphPad Prism 6.0 software. The MST data were analyzed using the NanoTemper analysis software (NanoTemper Technologies GmbH) and plotted using GraphPad Prism 6.0 software. Statistical parameters are reported in the Figure Legends.

### DATA AND SOFTWARE AVAILABILITY

The AcrIIA1 homolog protein accession numbers and associated promoter sequences are disclosed in Table S1.

